# Key Proteins for Regeneration in *A. mexicanum*: Transcriptomic Insights from Aged and Juvenile Limbs

**DOI:** 10.1101/2023.09.07.556684

**Authors:** Aylin del Moral-Morales, Cynthia Sámano, José Antonio Ocampo-Cervantes, Maya Topf, Jan Baumbach, Rodrigo González-Barrios, Ernesto Soto-Reyes

## Abstract

The axolotl is an animal with remarkable regenerative abilities, making it an ideal model for studying potential regenerative therapies in mammals, including humans. However, the molecular mechanisms involved in regeneration remain unclear. We conducted a transcriptomic analysis of juvenile axolotls’ limbs and their blastema and compared the results with aged axolotls that failed to regenerate after amputation. We identified a set of genes involved in cell differentiation, transcriptional regulation, cartilage development, bone morphogenesis, and extracellular matrix remodeling. Four highly expressed genes (*FSTL1, ADAMTS17, GPX7*, and *CTHRC1*) were identified in regenerating tissue, but underexpressed in aged axolotls. Structural and homology analysis showed that these genes are conserved and have important roles in development, bone morphogenesis, and cartilage formation. Our findings propose a novel set of axolotl genes involved in tissue regeneration that could be a starting point for further studies in other vertebrates.

**Graphical abstract:** 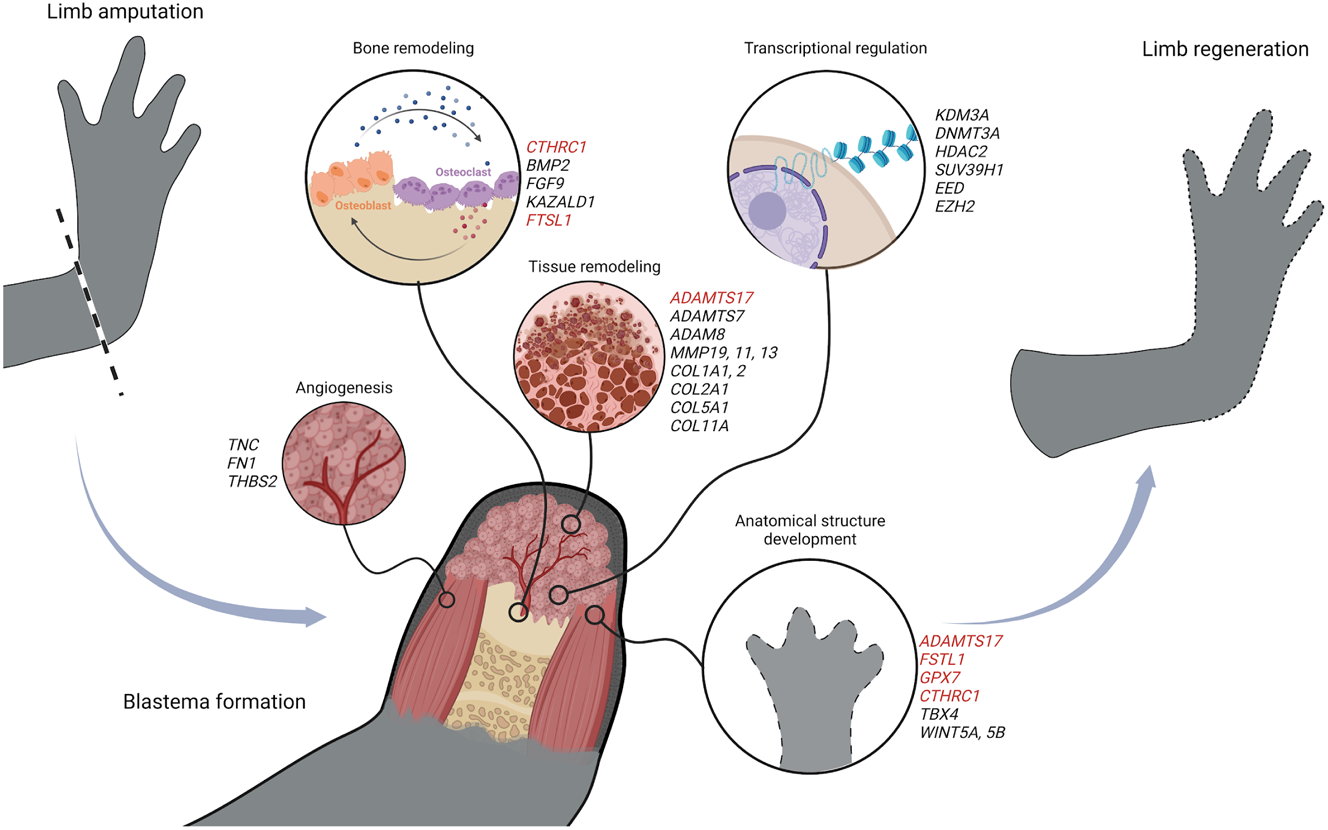

## Introduction

The axolotl is a salamander of the genus *Ambystoma*, which has a distribution of 32 species from southern Canada to central Mexico. Among the different species present in Mexico, *Ambystoma mexicanum*, an endemic species from the lake region of Mexico City, is the most studied (Sámano et al., 2021; Vance, 2017). Commonly known as axolotl, *A. mexicanum* is a vertebrate amphibian with an unusual appearance. Unlike other salamanders, axolotls do not metamorphose to become adults, but instead remain in their larval form, retaining their external gills and aquatic lifestyle throughout their entire lives (Demircan et al., 2016; Monaghan et al., 2012). In addition to their popularity as pets in many parts of the world, this animal is considered an interesting model for studying tissue regeneration. Indeed, they can regenerate a wide range of injured tissues in just a few weeks, including limbs (Haas & Whited, 2017; Simon & Tanaka, 2013), tail, central and peripheral nervous system (McHedlishvili et al., 2012; Tazaki et al., 2017), iris (Suetsugu-Maki et al., 2012), bone and muscle (Joven et al., 2019; Wu et al., 2015).

According to previous studies, the axolotl regeneration process involves 3 phases: wound healing, blastema formation, and re-development (Knapp et al., 2013; Stocum & Cameron, 2011). After an injury, the fibroblasts surrounding the wound acquire proliferative capacity and set the stage for new tissue development (S. V. Bryant et al., 2004; Lin et al., 2021; Masselink & Tanaka, 2021; Sandoval-Guzmán et al., 2014). Once the wound is closed, the fibroblasts dedifferentiate into mesenchymal progenitor cells that accumulate in an epithelium-covered region called the blastema. It is in this organ where the reconstruction of the missing limb takes place. Blastema cells are thought to have a certain “cellular memory”, as they always differentiate into the same cell type from which they originated, and the new limb will have the same size, shape, and orientation as the amputated one (Kragl et al., 2009; McCusker et al., 2015; McCusker & Gardiner, 2011). Because of the former, the study of the underlying mechanisms and factors involved in blastema regeneration is currently of great interest. Specifically, researchers are interested in dissecting the cellular and molecular events leading to regeneration in axolotls that could be extrapolated to other species. It was shown in a previous study that mouse growth factors can induce blastema formation in axolotls, suggesting that the machinery required for tissue regeneration may be conserved in mammals (Makanae et al., 2014). However, the molecular factors involved in the activation of such machinery remain unexplored.

One limitation of studying axolotls is the complexity of their genome; with 32 gigabase pairs of DNA distributed across 14 pairs of chromosomes, it is one of the largest genomes among vertebrates. However, approximately 70% of their genome is composed of repetitive elements, making its assembly a challenge (Keinath et al., 2015). Furthermore, the lack of a fully annotated transcriptome has hampered comprehensive studies of tissue regeneration, making it difficult to compare results across publications. Despite these challenges, previous research has identified several candidate genes that may play a role in the regenerative process. However, the mechanistic insights into these genes and how they interact with others are not fully understood (D. M. Bryant et al., 2017; Caballero-Pérez et al., 2018; Darnet et al., 2019). Fortunately, recent advances in genome assembly and transcriptome annotation are enabling detailed transcriptomic studies that will shed light on the molecular processes underlying axolotl regeneration (Schloissnig et al., 2021).

Furthermore, the impact of environmental factors, such as habitat, diet, and age, on the regenerative capacity of axolotls has not been investigated (Haas & Whited, 2017). While axolotls can regenerate tissues throughout most of their entire lifespan, this phenomenon becomes less efficient with age (Sousounis et al., 2014; Suetsugu-Maki et al., 2012). Younger individuals can regenerate tissues in a matter of days or weeks, whereas sexually mature adults may require several months to regenerate a limb (McCusker & Gardiner, 2011; Vieira et al., 2020). It is also worth noting that most experiments have been carried out on the *d/d* strain, a homozygous white mutant lineage established by French naturalists in the mid-19th century. Interestingly, this strain often exhibits regeneration defects due to extensive inbreeding (Farkas & Monaghan, 2015; Sámano et al., 2021). Therefore, to fully understand the genetic, transcriptomic and environmental factors involved in tissue regeneration in *A. mexicanum*, it is crucial to study native Mexican axolotl populations from the Xochimilco Lake area.

In this study, we analyzed transcriptomic data from the limbs of juvenile axolotls (eight months old) and the blastema formed 10 days after amputation to investigate the molecular signatures associated with tissue regeneration. Additionally, we collected samples from the posterior leg of two aged axolotls (eight years old) that did not develop a blastema after amputation. By comparing our datasets with those previously reported by Bryant et al. in 2017 (D. M. Bryant et al., 2017) we aimed to identify common transcriptomic responses in blastemas. Through a custom annotation of the axolotl proteome, we found that the differentially expressed genes were primarily associated with anatomical development and cell differentiation. Among these, we identified four key genes (*FSTL1, ADAMTS17, GPX7*, and *CTHRC1*) that were highly expressed in regenerating tissues but were underexpressed in aged axolotls. These genes showed association with other genes involved in anatomical structure development. Structural and conservation analyses revealed that these proteins are highly conserved in vertebrates and play important roles in development, bone morphogenesis, and cartilage formation in mammals. Our results suggest a set of key axolotl genes that potentially participate in tissue regeneration specifically in juvenile axolotls. These genes could provide a basis for studying tissue regeneration in other vertebrates and they may have implications for the field of regenerative medicine in the future.

## Results

### Blastemas share a defined transcriptomic profile

In order to study the genes that are expressed during the formation of the blastema in *A. mexicanum*, we collected RNA samples from the lower limbs of five juvenile axolotls (eight months old) endemic to Lake Xochimilco in Mexico City. The blastema sample was collected ten days after amputation and RNA was extracted. The samples were then analyzed by sequencing. We compared our results with RNA-seq data from Bryant et al. (D. M. Bryant et al., 2017), obtained from strain *d/d* axolotls (white mutant) that had their forelimbs amputated at two different sites. To identify the most significant changes in gene expression between the different samples, we performed differential expression analysis (**Fig 1A**). Overall, we observed that most of the differentially expressed genes (DEGs) were downregulated. In our samples, we identified 667 upregulated genes and 2076 downregulated genes in the blastema compared to the control tissue. In the distal blastema, we found 4809 upregulated genes and 5577 downregulated genes compared to the control, whereas in the proximal blastema, we found 6143 upregulated genes and 7671 downregulated genes (**Fig 1B**).

**Figure 1.**
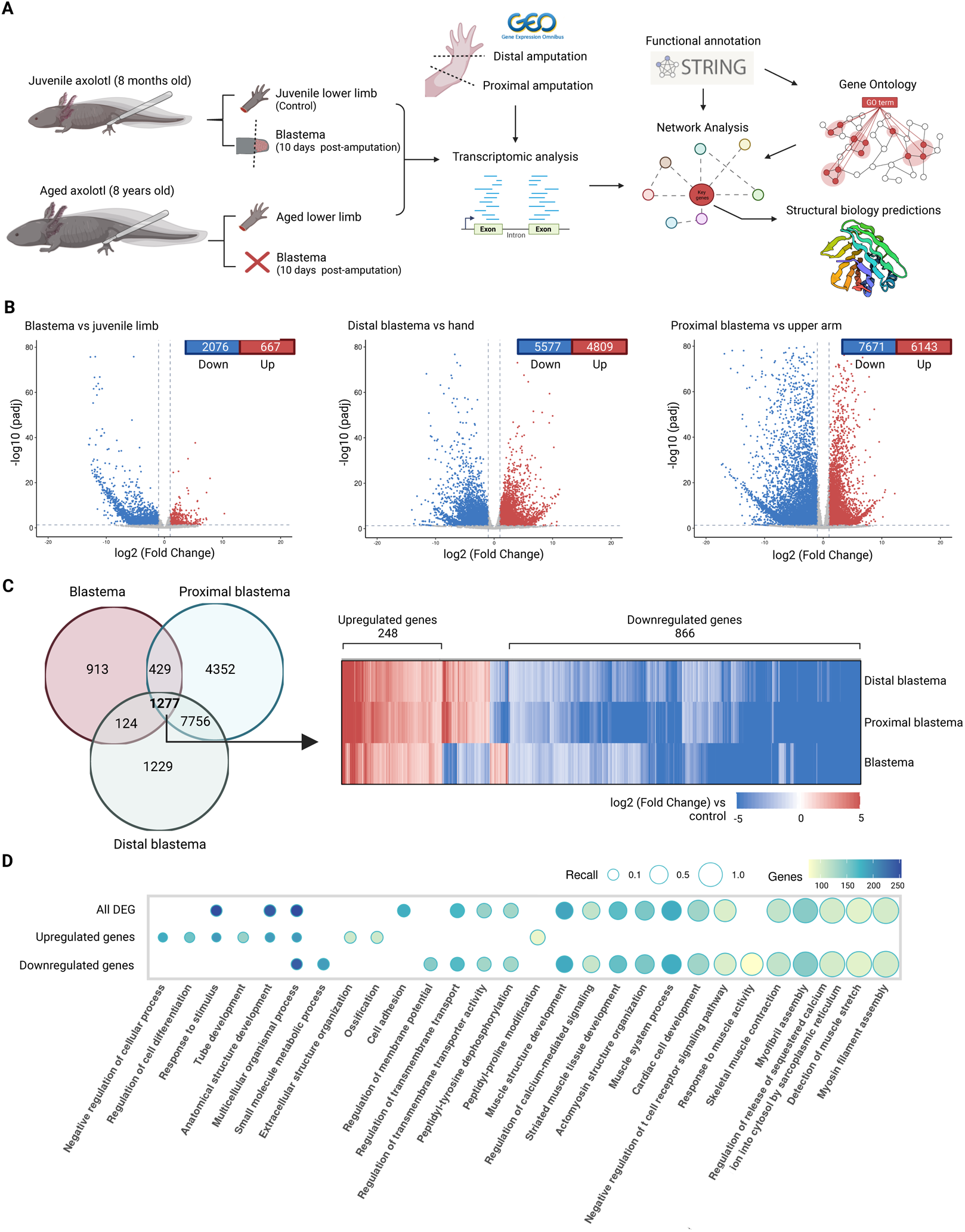
Comparison of DEG in different blastema samples. A Schematic representation of tissue sampling and downstream analyses. We sampled 5 juvenile axolotl limbs (control), 5 blastemas, and 2 aged axolotl limbs (over eight years old). Data sets published by Bryant et al. were also analyzed. Samples were taken from two different amputation sites on the axolotl forelimbs. The first was at hand level (distal blastema) and the second was below the elbow (proximal blastema). After RNA extraction and sequencing, differential expression and network analysis were performed to find relevant genes and predicted GO terms associated with regeneration. B Differentially expressed gene (DEG) volcano plots. The left panel shows the comparison between blastema and control limbs (juvenile axolotls). The middle panel shows distal blastema compared to the hand. The right panel shows proximal blastema compared to the upper arm. Red dots represent upregulated genes (p-value < 0.05 and log2(Fold Change) > 1). Blue dots represent the downregulated genes (p-value < 0.05 and log2(Fold Change) > 1). C Venn diagram of the intersection between DEG for blastema vs. control, distal blastema vs. hand, and proximal blastema vs. humerus. The left panel shows the heat map for the 1277 genes in the intersection between DEG in aged axolotls and blastema. The color of the tiles represents the log2(Fold Change) value for each gene. Genes with the same behavior across samples are indicated by parentheses (1134 of 1277 genes). D Top 20 Gene Ontology biological processes associated with all DEG in the intersection shown in Fig 1C (1134 genes), downregulated genes only (886 genes), or upregulated genes (248 genes). The size of the bar corresponds to the number of genes associated with the significant GO term. The color of the bars represents the log10(p-value) of the term. Created with BioRender.

We compared the differentially expressed genes (DEGs) in our samples with Bryant’s datasets to identify genes that were consistently regulated across experiments. We found 1277 DEGs that were common to all three datasets. Specifically, out of these common DEGs, 248 were upregulated and 886 were downregulated in all three samples (**Fig 1C, Dataset EV1**). Since ontology annotation for *A. mexicanum* is not available, we used STRING (Szklarczyk et al., 2023) to generate a homology annotation for the coding genes in the *AmexT_v47* transcriptome. By performing a gene ontology term enrichment analysis, we found that the DEGs in the blastema were primarily associated with biological processes related to tissue and muscle development, cytoskeleton organization, cell adhesion, extracellular matrix organization, and ossification. However, among the downregulated genes, we observed an enrichment in biological processes such as cytoskeleton organization, muscle structure development, and myofibril assembly. Conversely, the upregulated genes were enriched in terms such as anatomical structure development, negative regulation of cellular processes, and regulation of cell differentiation (**Fig 1D, Dataset EV2**).

A gene-GO term network was constructed using ten of the most representative terms from **Fig 1D** to provide a more comprehensive visualization of the genes involved in the process of regeneration. Notably, the majority of genes belonged to the term anatomical structure development, and a significant proportion of the genes in this category showed downregulation in blastema compared to control. However, genes associated with the Wnt pathway (*WNT5A* and *WNT5B*) were upregulated, as well as the metallopeptidases *ADAMTS7*, *ADAM8*, *MMP19*, *MMP11*, and *MMP13*, which are also associated with the extracellular structure organization term. Similarly, genes related to cell adhesion, myofibril assembly, and actomyosin structure organization were mostly downregulated in blastemas, with a few exceptions such as tenascin (*TNC*), fibronectin (*FN1*), and thrombospondin-2 (*THBS2*). Another group of genes, including *SMAD6*, *SMAD7*, and *BAMBI*, which are involved in the tumor growth factor beta (TGF-β) signaling pathway, were upregulated and associated with cell differentiation, along with *SOX4* and *NOX4*. Genes associated with epigenetic functions were also found. A notable example is the histone lysine demethylase *KDM3A,* which was found among the genes involved in differentiation and development. This finding suggests tissue regeneration may be controlled by epigenetic processes. Notably, genes involved in ossification and regulation of cell differentiation were also upregulated. Additionally, several genes previously associated with regeneration in axolotls, marked in squared boxes, were obtained from the table published by Haas and Whited (Haas & Whited, 2017). Among these genes, *KAZALD1* stands out as a well-studied gene involved in anatomical development and bone regeneration in axolotls (D. M. Bryant et al., 2017). Furthermore, the Bone Morphogenetic Protein 2 (*BMP2*) and the Fibroblast Growth Factor 9 (*FGF9*), which are associated with skeletal and cartilage development, are upregulated. These proteins are particularly relevant because a previous study demonstrated their ability to induce blastema formation in axolotls (Makanae et al., 2014). To summarize, our results provide a collection of genes and biological processes that are associated with the blastema tissue and the regeneration process in axolotls, which is consistent with and extends the data from previous studies (**Fig 2**).

**Figure 2.**
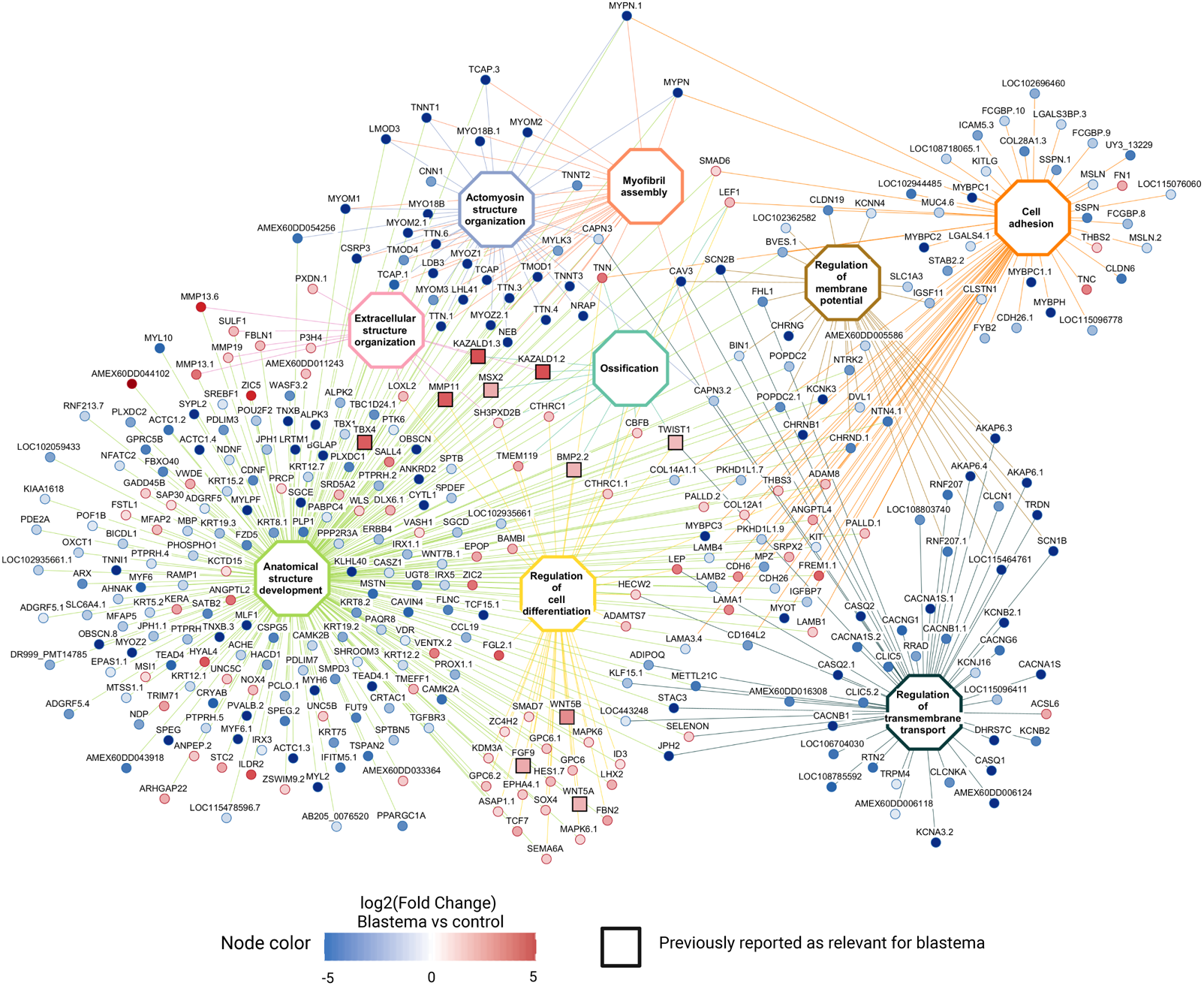
Gene Ontology terms and genes associated with axolotl regeneration. Gene Ontology terms enriched for the DEG in the three blastema datasets analyzed. Node color corresponds to the log2(Fold Change) for the blastema vs. control dataset. GO terms are white and each one has a different border color. Edges are colored according to the GO term from which they originate from. Squared nodes represent genes that have been previously reported in the literature to be associated with the axolotl regenerative process. Created with BioRender.

### Key genes in regeneration revealed by the transcriptomic profile of aged axolotls

In addition to the previously collected samples, two aged axolotls (eight years old) were subjected to limb amputation; however, they showed no signs of regeneration after 10 days, in contrast to the juvenile axolotls; thus, no post-amputation sample was collected for this group. When we compared the transcriptome of the aged vs. juvenile limbs, only 172 genes were differentially expressed, and most were downregulated compared to the control (**Fig 3A**). After a GO enrichment analysis, the only significant GO term for the DEG in aged limbs was collagen fibril organization. To search for relationships among the aged DEGs, *de novo* pathway enrichment was performed with KeyPathwayMineR (Mechteridis et al., 2022). As input, we used the DEGs in aged limbs and a protein-protein interaction network constructed with STRING using the proteome from the *AmexT_v47* annotation file. The interaction network shows mainly the downregulation of type I (*COL1A1, COL1A2*), II (*COL2A1*), V (*COL5A1*), and XI (*COL11A1*) collagens. Some of these proteins appear more than once because they were assigned the same name, according to the transcriptome annotation file, so they were marked with a numerical suffix. There are also several ribosomal components, such as the ribosomal proteins RPS2 and RPS5, as well as the signal recognition particle 9 *(SRP9*) and *SEC61G*, which are part of the complex required for protein translation at the endoplasmic reticulum (**Fig 3B**).

**Figure 3.**
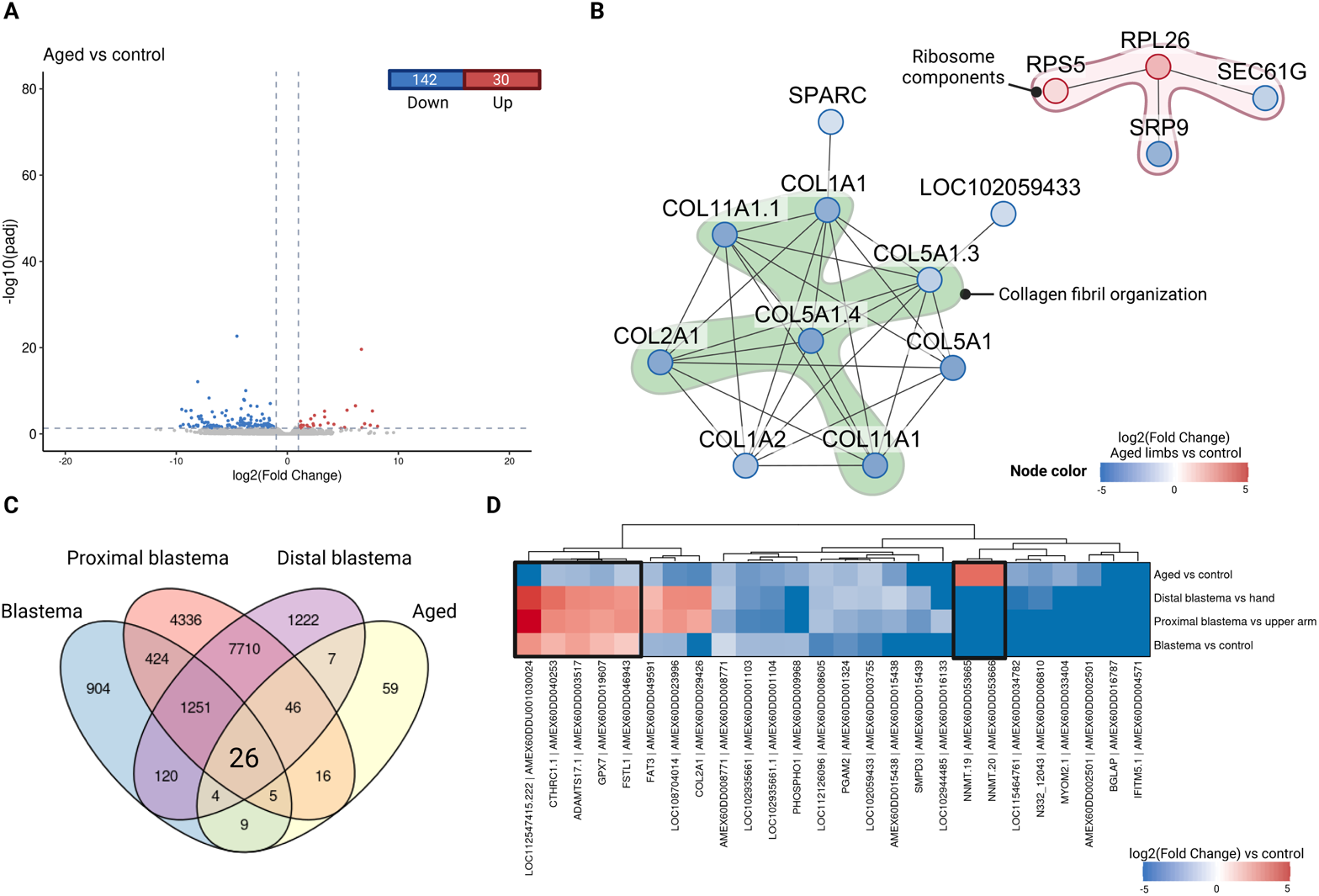
Differential Expressed Genes in aged axolotls. A Volcano plot of the differentially expressed genes (DEGs) obtained for the comparison between aged limbs vs. control limbs (juvenile axolotls). Red dots represent the upregulated genes (p-value < 0.05 and log2 (Fold Change) > 1). Blue dots represent the downregulated genes (p-value < 0.05 and log2 (Fold Change) < - 1). B Protein-protein interaction network for the DEGs in aged limbs. Node color represents the log2 (Fold Change) of aged limbs vs. control. Genes associated with the term collagen fibril organization are highlighted in green. Genes that are part of the ribosome are highlighted in pink. Edges represent predicted interactions between the proteins encoded by the genes. C Venn diagram for the DEGs in all four datasets evaluated. D Heat map with the fold change of all the DEGs in the intersection shown in panel c (26 genes). DEGs that show contrasting patterns in the blastema vs. aged datasets are highlighted with a black box. Created with BioRender.

In an effort to identify key genes involved in the tissue regeneration process, we focused on the differentially expressed genes (DEGs) that showed contrasting patterns in the blastema vs. control and aged vs. control comparisons. Our goal was to identify genes that were upregulated in blastemas but downregulated in old axolotls, and vice versa, as they could offer valuable insights into the impaired tissue regeneration observed in aging axolotls. Through DEG overlap analysis, we discovered 26 genes that consistently exhibited significant differential expression across all four datasets evaluated (**Fig 3C**). Among these genes, 7 showed consistent differential expression specifically in blastemas compared to aged limbs. Notably, *CTHRC1.1*, *ADAMTS17.1*, *GPX7*, *FSTL1*, and *LOC112547415.222* were found to be upregulated in blastemas but downregulated in aged axolotls. In contrast, *NNMT.19* and *NNMT.20* showed the opposite pattern, being upregulated in aged axolotls but downregulated in blastemas (**Fig 3D**). These genes hold significant interest because their high expression in blastemas and downregulation in aged axolotls, which have lost their regenerative capacity, suggests that they may play a critical role in the process of regeneration. Therefore, we will refer to this set of genes as regeneration-related genes. **Table 1** summarizes the annotation and putative function of the regeneration-related genes found.

**Table 1.** Annotation and putative function of the regeneration-related genes found. Gene ID is according to the *AmexT_v47* transcriptome. The homolog sequence is the same as provided in the file, which is the closest homolog in the NCBI reference database.

| Gene ID | Gene Name | Homolog | Location |  |  |
| --- | --- | --- | --- | --- | --- |
|  |  |  | Chromosome | Start | End |
| AMEX60DD003517 | *ADAMTS17.1* | *ADAMTS17*<br>A disintegrin and metalloproteinase with<br>thrombospondin motifs 17 *Terrapene carolina*<br>*triunguis*<br>NCBI Reference Sequence: XP_026504145.1 | chr11p | 81488489 | 82602596 |
| AMEX60DD019607 | *GPX7* | *GPX7*<br>PREDICTED: glutathione peroxidase 7<br>*Latimeria chalumnae*<br>NCBI Reference Sequence: XP_006006032.1 | chr1q | 1276741180 | 1277033638 |
| AMEX60DD040253 | *CTHRC1.1* | *CTHRC1*<br>collagen triple helix repeat-containing protein 1<br>*Xenopus laevis*<br>NCBI Reference Sequence: XP_018123683.1 | chr5q | 899669621 | 899687475 |
| AMEX60DD046943 | *FSTL1* | *FSTL1*<br>follistatin like 1<br>*Rhinatrema bivittatum*<br>NCBI Reference Sequence: XP_026504145.1 | chr7p | 70865121 | 71016956 |
| AMEX60DDU001030024 | *LOC112547415.222* | uncharacterized protein LOC112547415<br>*Pelodiscus sinensis*<br>NCBI Reference Sequence:<br>XP_025045344.1 | C0173161 | 1 | 8438 |
| AMEX60DD053665 | *NNMT.19* | *NNMT*<br>Indolethylamine N-methyltransferase-like<br>*Microcaecilia unicolor*<br>NCBI Reference Sequence: XP_030077372.1 | chr9p | 79243911 | 79434013 |
| AMEX60DD053666 | *NNMT.20* | *NNMT*<br>Nicotinamide N-methyltransferase-like<br>*Rhinatrema bivittatum*<br>NCBI Reference Sequence: XP_029428629.1 | chr9p | 79380383 | 79403217 |

### Co-expression network analysis reveals gene modules associated with tissue regeneration in Ambystoma mexicanum

As an additional approach, we used co-expression network analysis to examine sets of genes that might be involved in regeneration and show similar transcriptional responses to our set of regeneration-related genes. This analysis incorporated muscle, cartilage, and bone samples from the study by Bryant et al. (D. M. Bryant et al., 2017), as well as two leg samples from Caballero-Pérez et al (Caballero-Pérez et al., 2018). Inclusion of these datasets allowed for a more comprehensive and robust understanding of the genes that are co-regulated, and thus similarly responding, in *Ambystoma mexicanum*. The resulting network comprised 35 gene modules, out of which 18 modules showed significant associations (p-value < 0.05 and absolute Pearson correlation coefficient > 0.5) with the evaluated conditions (**Appendix Figure S1**, **Dataset EV3**). Four of the significant modules showed a strong correlation with the blastema condition (yellow, magenta, darkorange, and black). The yellow and magenta modules were also significantly associated with muscle samples. Genes involved in metabolic processes and muscle structure development are enriched in the yellow module, while the magenta module showed enrichment in genes associated with the cell cycle, DNA replication, and RNA splicing. The black module is exclusively associated with blastema and was enriched in genes associated with developmental processes and multicellular organism development. Notably, the darkorange module did not show any significant gene ontology term association but is also associated with limb samples (**Fig 4A, Appendix Figure S2**).

**Figure 4.**
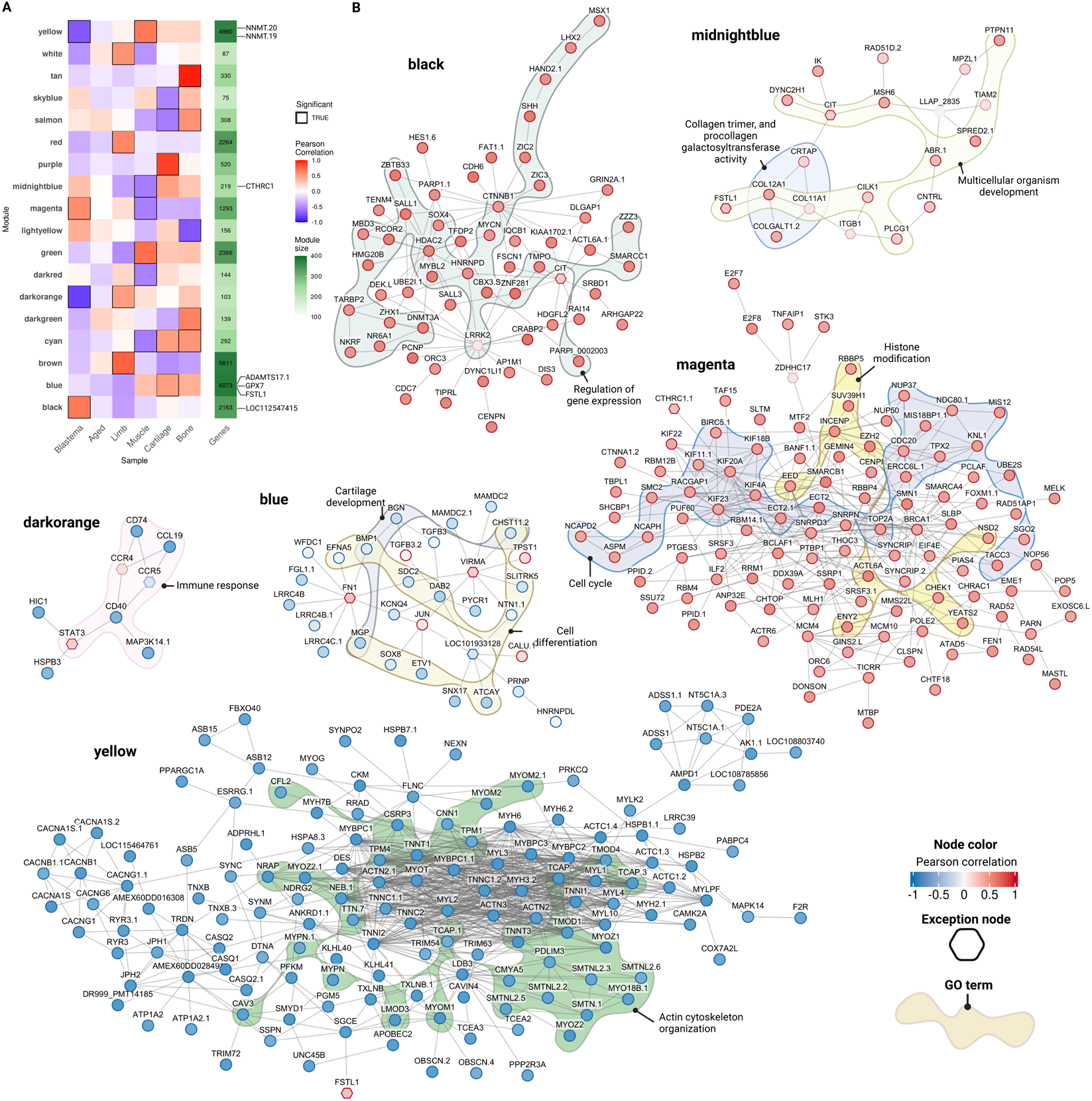
Gene coexpression analysis of axolotl samples. A Module-trait correlation for the 18 significant co-expression modules identified. Tile color represents the module-trait Pearson correlation coefficient. Significant modules (p-value < 0.05) are enclosed in a black box The green panel indicates the module size. B Overview of the co-expressed genes within the black, midnightblue, yellow, blue, darkorange, and magenta modules. Node color corresponds to the Pearson correlation coefficient for the blastema samples. Edges represent putative protein-protein interactions predicted by STRING. Gene Ontology terms associated with some nodes are highlighted with colored shapes. Exception nodes (genes not belonging to the module) are marked with a hexagon. Created with BioRender.

Although none of the modules showed a strong association with the aged samples, we identified a set of 19 genes that had a significant association (absolute module-trait correlation > 0.7 and p-value < 0.05) with the trait (**Dataset EV4**). Among these genes, the two with the strongest correlation were *AMEX60DD027179*, an non-annotated gene encoding a 127 amino acid protein of unknown function, and *PAFAH2*, a gene encoding platelet-activating factor acetylhydrolase isoform 2. Both genes were negatively associated with aged axolotls and belonged to the turquoise module, which exhibited enrichment in genes associated with the regulation of metabolic processes, organelle organization, and chromatin organization.

To gain further insight into the biological processes or pathways associated with each module, we used KeyPathwayMiner to construct protein-protein interaction (PPI) networks. **Fig 4B** displays the modules that were significant for blastema and those that contain one or more of the previously identified regeneration-related genes. Network nodes are color-coded based on their gene-trait correlation for blastema, while the edges represent predicted interactions between the proteins encoded by the genes. The colored shapes indicate gene ontology terms associated with the module nodes. *CTHRC1,* one of the regeneration-related genes, was found in the midnightblue module, which exhibited significance in muscle samples but had no enriched gene ontology terms. The network revealed proteins associated with multicellular development and collagen trimer. The blue module contained three out of the seven regeneration-related genes identified through differential expression analysis (*ADAMTS17.1*, *GPX7*, and *FSTL1*). This module displayed a strong correlation with cartilage samples and enrichment in genes associated with the regulation of developmental processes. The PPI network for this module showed enrichment of genes related to cartilage development (e.g., *BMP1* and *CHST11*) and cell differentiation (e.g., *SOX8*, *SDC2*, and *DAB2*). Furthermore, the dark orange module, which negatively correlated with blastema, showed enrichment in immune response-related proteins, as well as some members’ association with *STAT3*. The magenta module included genes related to the cell cycle, particularly belonging to the kinesins family (*KIF)*. Notably, genes related to histone modification were also observed. These included the Polycomb group members *EED* and *EZH2*, as well as *SUV39H1*, a histone lysine methyltransferase. Genes involved in the regulation of gene expression, including *HDAC2* (histone deacetylase), *DNMT3A* (DNA methyltransferase), and transcription factors such as *SOX4*, *MYCN*, and *SALL1*, were found in the PPI network for the black module, one of the modules closely associated with the blastema and containing the regeneration-related gene *LOC112547415.* Finally, the two *NNMT* genes were found in the yellow module, which was enriched with several proteins associated with the organization of the actin cytoskeleton, as well as various components of the cytoskeleton, such as troponin genes (*TNT*), actin (*ACTN*), myotilin (*MYOT*), and myosin (*MYL*). Interaction between *SGCE*, another protein from the cytoskeleton, and *FSTL1* was also observed in this module.

In summary, we identified gene modules associated with regeneration and genes with similar transcriptional responses to our set of regeneration-related genes. Notably, the identified modules included our proposed regeneration-related genes, as well as other genes with potential involvement in regeneration. These results highlight the potential interplay between the regeneration-related genes we identified and other factors involved in axolotl limb regeneration, as well as providing clues as to the potential biological significance of the identified genes.

### The regeneration-related genes are conserved in vertebrates

Through transcriptomic analysis, we have identified a group of genes relevant for regeneration in *A. mexicanum*. To assess the functionality and presence of the proteins encoded by these genes in other organisms, we performed a homology search against the UniProtKB + Swiss-Prot database (UniProt Consortium, 2023) using a selection of animals based on S. Hedges’ comprehensive review of model organisms (Hedges, 2002). **Fig 5** illustrates the identity between the proteins encoded by the regeneration-related genes and their closest homologues in each selected organism. The UniProt IDs for each sequence and the full set of results are given in **Dataset EV5.** It is noteworthy that all of the regeneration-related proteins in *A. mexicanum* show homology in vertebrates, although their presence in invertebrates is variable. Among the regeneration-related proteins, CTHRC1 has the highest percentage of identity in vertebrates, while the NNMT proteins have lower conservation and a lower percentage of positives, indicating that only a fraction of the target protein matches the query sequences. However, FSTL1 and ADAMTS17 also show conservation in vertebrates but not in invertebrates. GPX7 is the only protein from the regeneration-related genes that is found in all of the organisms evaluated, albeit with a noticeably lower percentage of identity in invertebrates compared to vertebrates. Notably, the gene *LOC112547415* does not appear in the graph as it lacks an associated open reading frame, leading us to hypothesize its potential as a non-coding RNA.

**Figure 5.**
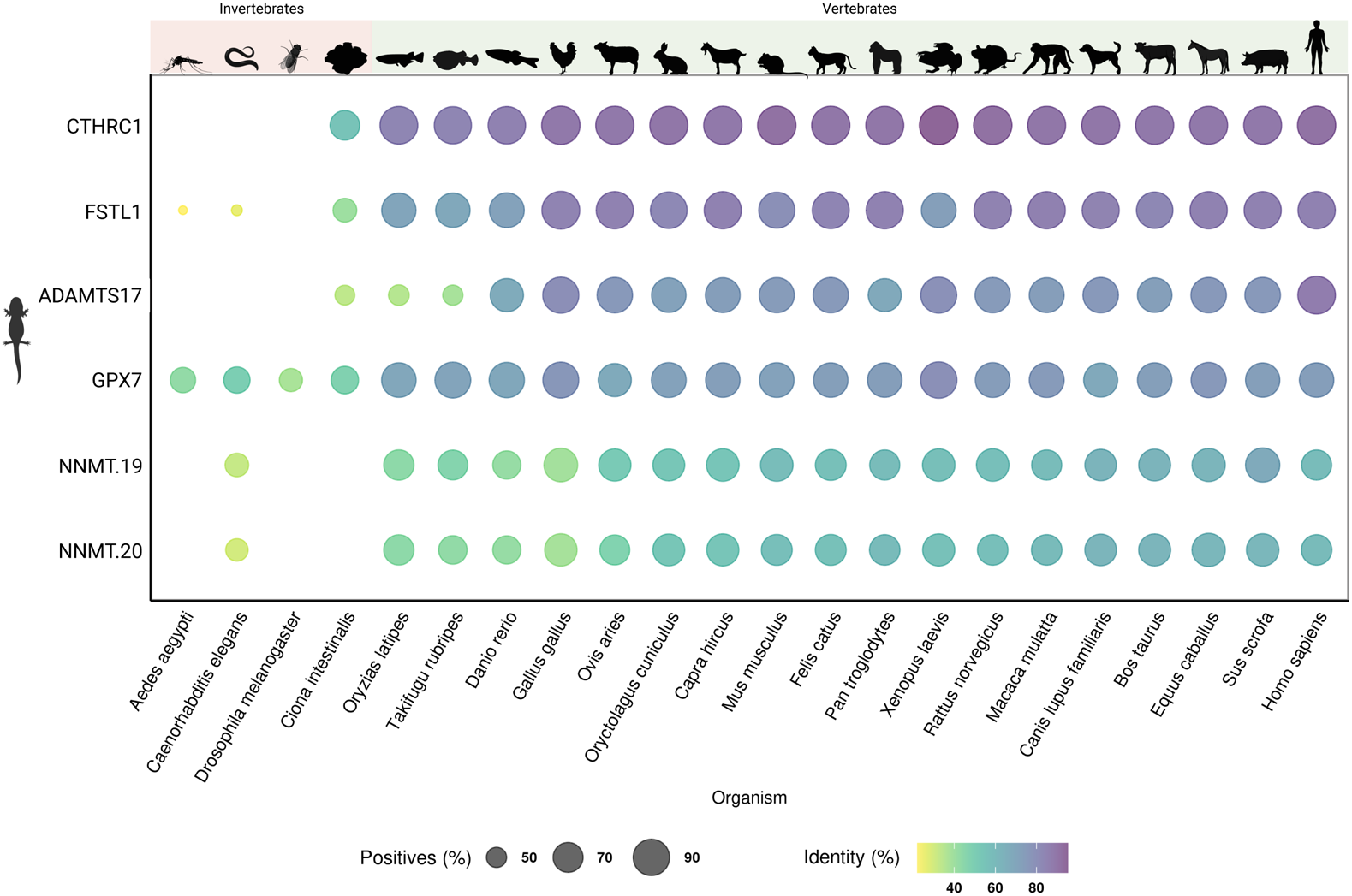
Conservation of the regeneration-related proteins in *A. mexicanum*. The plot shows the percentage of identity of each regeneration-related A. mexicanum protein (query) vs. its closest homolog in other selected animals (target). Bubble color indicates the percentage of identity between query and target protein. Bubble size is proportional to the percentage of positives. Positives are amino acids in the subject sequence that are identical or have similar chemical properties to the query sequence. Missing values indicate that no similar proteins were found in the subject organism. Created with BioRender.

Subsequently, the 3D structure of the proteins encoded by each regeneration-related gene in A. mexicanum was modeled using AlphaFold2 (Jumper et al., 2021), while sequence conservation was assessed using the Consurf (Yariv et al., 2023) server and the UniRef90 database. The complete predictions and confidence scores can be found in **Appendix Figure S3**. In addition, the Conserved Domain Database (Marchler-Bauer et al., 2015) was used to identify conserved domains within the proteins, with the details summarized in **Table 2**. Regarding the NNMT proteins, they share 97% identity with each other (**Appendix Figure S4**), and both have an S-adenosylmethionine-dependent methyltransferases domain (**Fig 6A**). NNMT.19 also possesses an extra N-terminal domain that does not appear to be conserved, as indicated by the “insufficient data” label from ConSurf (**Fig 6B**). However, upon comparing NNMT.19 with a previously published crystallographic structure of human NNMT (Y. Peng et al., 2011) it became evident that both axolotl proteins possessed an incomplete catalytic domain, suggesting that only half of the protein is present in axolotls (**Fig 6C)**, consistent with the low percentage of positive amino acids found previously.

**Figure 6.**
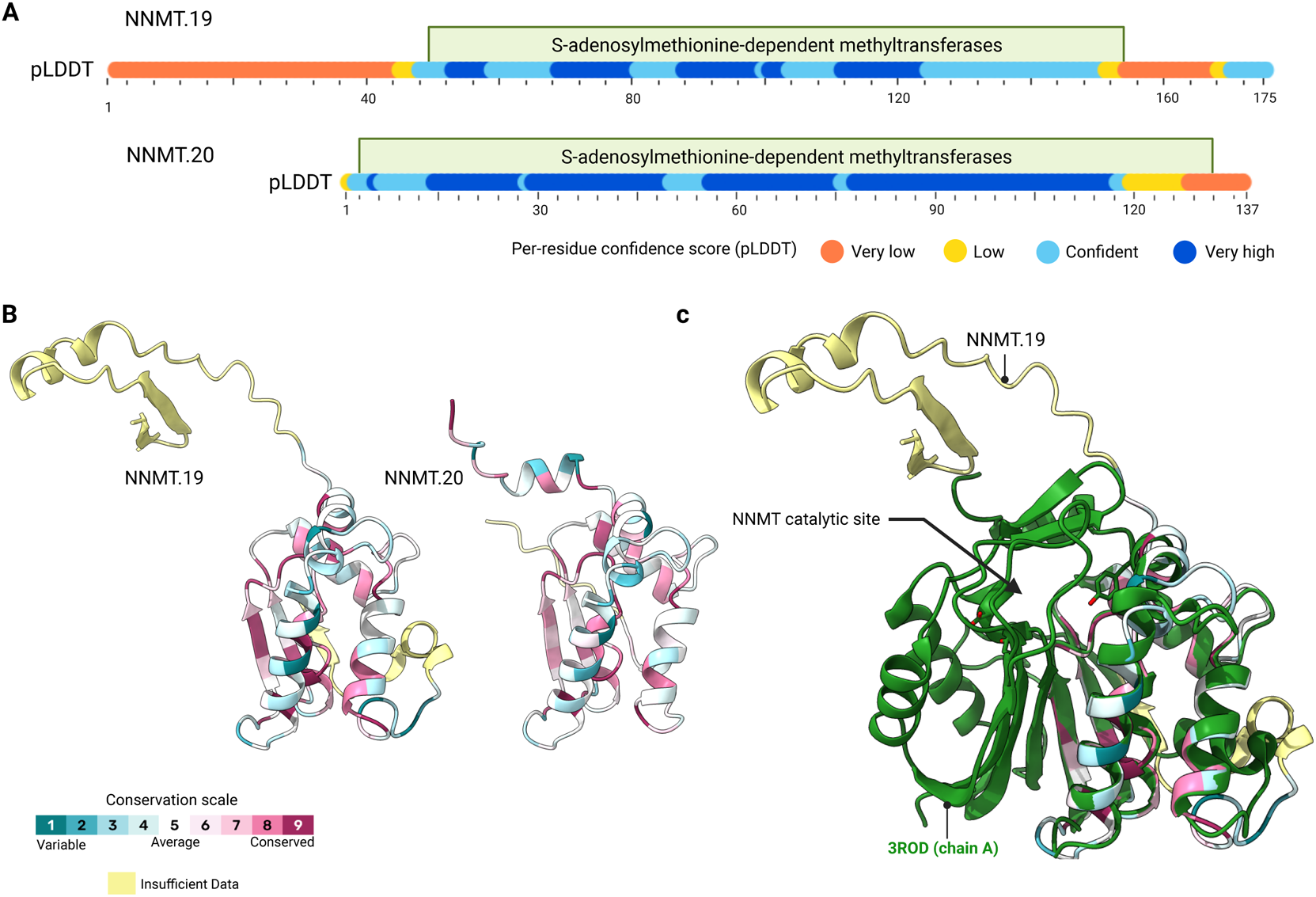
Sequence and conservation of *A. mexicanum* NNMT proteins. A Domain overview of A.mexicanum NNMT.19 and NNMT.20 proteins. The figures represent the sequence alignment of the proteins and the scale corresponds to the number of amino acids in each protein. The predicted conserved domains are annotated with a green box. The per residue confidence (pLDDT) for the AlphaFold models is represented with a bar. The complete structures can be found in Supplementary Fig 3. B AlphaFold2 structure prediction for both NNMT proteins. The predictions were obtained with the monomer preset for AlphaFold and using templates. The structures are colored according to their sequence conservation as calculated by the ConSurf server (ref). C Structural alignment between A. mexicanum NNMT.19 and an X-ray crystallography structure for human Nicotinamide N-methyltransferase (NNMT) deposited in PDB (ID: 3ROD). Created with BioRender.

**Table 2.** Domain annotation for the regeneration-related A. mexicanum proteins. The domain ID is from the Conserved Domain Database (CDD).

| Name | Gene ID | Transcript ID | E-Value | Residue Range | Domain |
| --- | --- | --- | --- | --- | --- |
| NNMT.20 | AMEX60DD053666 | AMEX60DD201053666.1 | 1.80e-48 | 5-133 | ID: cl17173<br>S-adenosylmethionine-dependent methyltransferases |
| NNMT.19 | AMEX60DD053665 | AMEX60DD301053665.1 | 8.28e-41 | 52-155 | ID: cl17173<br>S-adenosylmethionine-dependent methyltransferases |
| ADAMTS17 | AMEX60DD003517 | AMEX60DD201003517.1 | 1.18e-24 | 26-163 | ID: pfam01562<br>Reprolysin family propeptide |
|  |  |  | 3.56e-84 | 217-434 | ID: cd04273<br>Zinc-dependent metalloprotease, ADAMTS_like subgroup. |
|  |  |  | 7.95e-23 | 451-518 | ID: cl20316<br>ADAM cysteine-rich domain |
|  |  |  | 7.31e-16 | 531-583 | ID: smart00209<br>TSP-1 type 1 repeat |
|  |  |  | 1.26e-08 | 615-684 | ID: cl41950<br>ADAM cysteine-rich domain |
|  |  |  | 3.06e-07 | 821-873 | ID: pfam19030<br>TSP-1 type 1 doamin |
|  |  |  | 3.10e-10 | 937-982 | ID: pfam19030<br>TSP-1 type 1 doamin |
|  |  |  | 2.14e-12 | 991-1040 | ID: pfam19030<br>TSP-1 type 1 doamin |
| FSTL1 | AMEX60DD046943 | AMEX60DD201046943.1 | 3.46e-12 | 30-99 | ID: cd01328<br>Kazal-type serine protease domain |
|  |  |  | 4.27e-73 | 112-225 | ID: cd16233<br>EF-hand |
| GPX7 | AMEX60DD019607 | AMEX60DD301019607.1 | 5.40e-10<br>5 | 46-198 | ID: cl00388<br>Thioredoxin-like catalytic residues |

ADAMTS17 from *A. mexicanum* has several conserved domains, including a catalytic domain (Kavanagh et al., 2007) with the zinc-binding HExxHxxGxxH consensus motif, characteristic of the catalytic site of the ADAMTS family of metalloproteases (Jones & Riley, 2005; Porter et al., 2005; Tang, 2001). It also contains a propeptide from the reprolysin family, two cysteine-rich regions and several TSP-1 motifs. A comparison with the crystallographic structure of human ADAMTS5 (Mosyak et al., 2008), an enzyme of the same family, shows that the catalytic residues of the axolotl’s ADAMTS17 are present, and that the sequence around them is conserved, suggesting that this protein could be catalytically active (**Fig 7A**). On the other hand, two domains were identified for FSTL1; a Kazal-type serine protease inhibitor and an EF-hand domain. The ConSurf analysis shows that FSTL1 from *A. mexicanum* is highly conserved. We particularly show a set of disulfide bonds with high conservation that together with the Kasal domain corresponds to the follistatin-like 1 (Fstl1-FK) domain described by Li et al (Li et al., 2019) (**Fig 7B**). GPX7 has only one thioredoxin-like domain; however, through the structural analyses, we found that the catalytic site and catalytic residues of the protein are conserved in *A. mexicanum*. Here we show a comparison between the catalytic site of *H. sapiens* GPX7 (Kavanagh et al., 2007) and the predicted structure for *A. mexicanum* GPX7; the residues W164, C79 and Q114, which are required for GPX7 activity, are also present in our prediction (**Fig 7C**).

**Figure 7.**
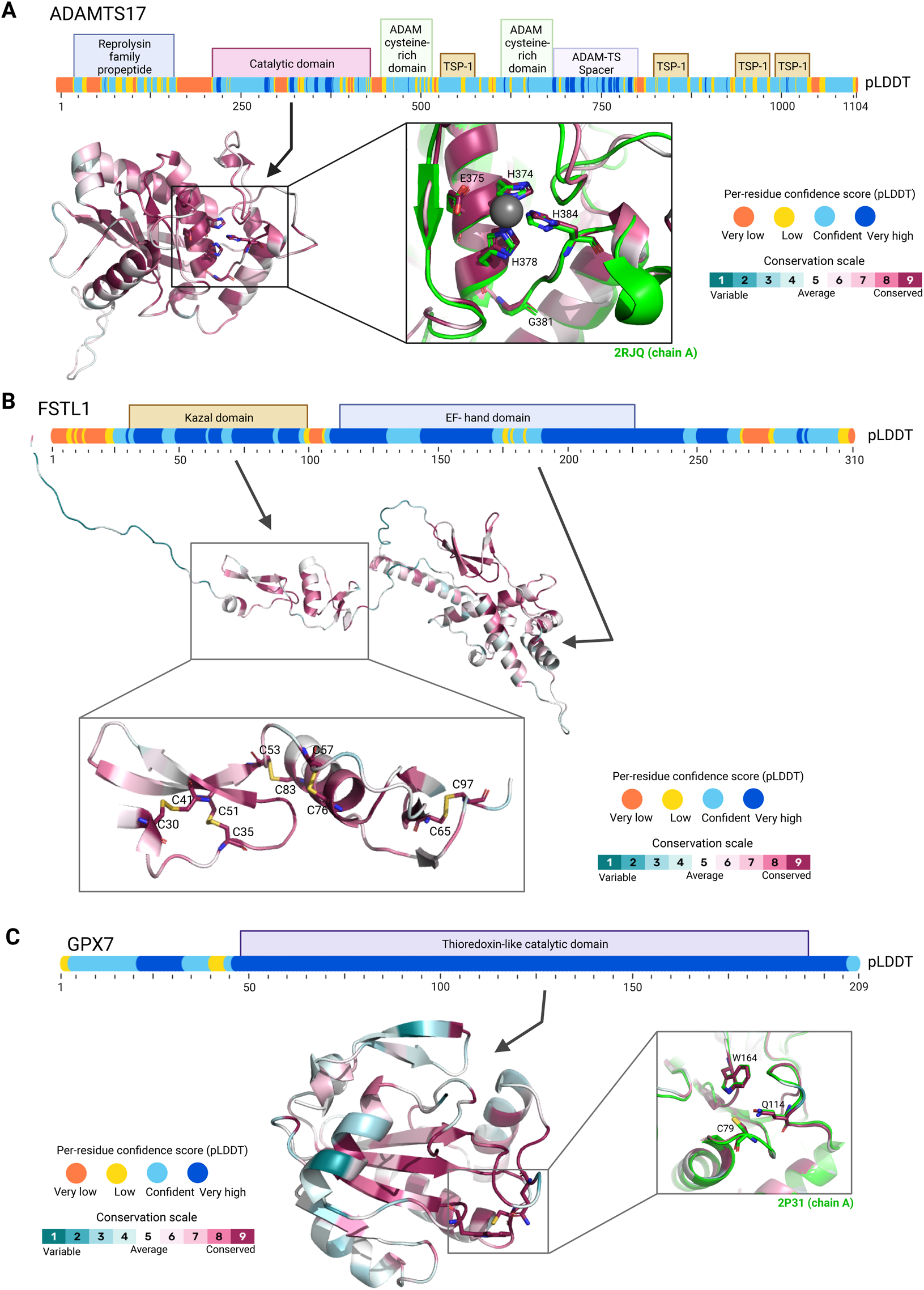
Conservation of ADAMTS17, FSTL1 and GPX7 proteins in *A. mexicanum*. Structures were predicted with AlphaFold2 and the conservation analysis was performed with the ConSurf server. A Domain overview and predicted structure of the catalytic domain of A.mexicanum ADAMTS17. Residues are colored according to their degree of conservation. The box shows the structural alignment of the catalytic sites of A.mexicanum ADAMTS17 and H. sapiens ADAMTS5 (PDB: 2RJQ, green). The per residue confidence (pLDDT) for the AlphaFold models is represented with a bar. The complete structures can be found in Supplementary Fig 3. B Predicted domains and structure of A.mexicanum FSTL1. Residues are colored according to their degree of conservation. The box shows the conservation of the cysteine bonds required for FSTL1 folding. The colored bar represents the per residue confidence (pLDDT) for the AlphaFold model. C Domain overview and predicted structure of the catalytic domain of A.mexicanum GPX7. The residues are colored according to their degree of conservation. The box shows the structural alignment of the catalytic sites of A. mexicanum GPX7 and H. sapiens GPX7 (PDB: 2P31, green). The colored bar represents the per residue confidence (pLDDT) for the AlphaFold model. Created with BioRender.

Finally, no conserved domain was identified for the CTHRC1 protein; however, structural prediction and conservation analysis suggest its conservation (**Fig 8A**). CTHRC1, also known as collagen triple helix repeat containing-1, is a secreted protein involved in osteogenesis and bone remodeling processes and typically exists as a homotrimer (Leclère et al., 2020). Using the AlphaFold multimer model, we obtained a high confidence structure for the axolotl CTHRC1 trimmer (**Fig 8B**). Taken together, our results suggest that the regeneration-related genes highlighted in this study code for proteins that are conserved in other vertebrates, allowing us to infer their function based on their homologues.

**Figure 8.**
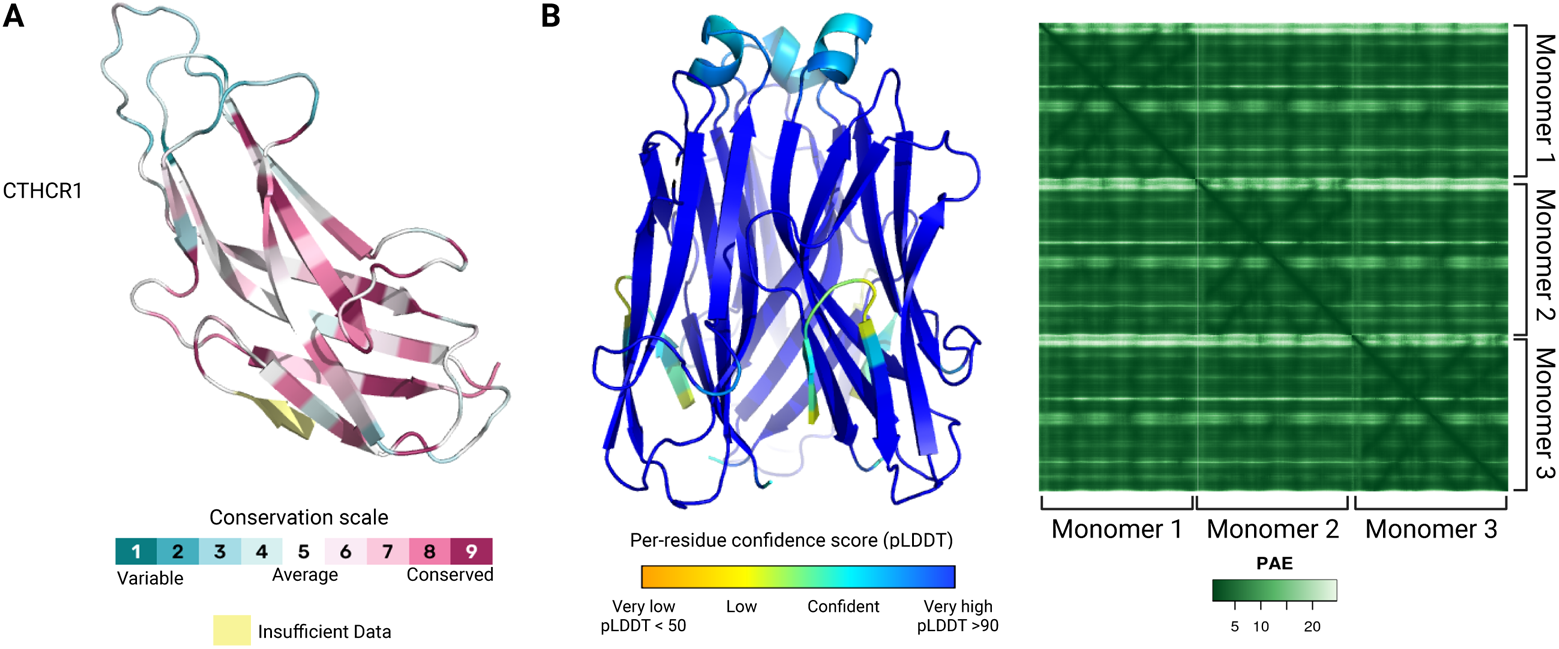
CTHRC1 conservation in A. mexicanum. A Complete predicted structure for A. mexicanum CTHRC1. The 3D structure prediction was obtained using AlphaFold2. Residues are colored according to their degree of conservation as calculated by the ConSurf server. B Heterotrimer 3D model prediction for CTHRC1. The prediction was obtained with AlphaFold2 multimer and the residues are colored according to the per-residue confidence score. The right panel shows the Predicted Aligned Error for the trimer. Low values mean higher confidence in the residue-residue interaction. Created with BioRender.

## Discussion

The *Ambystoma mexicanum*, commonly known as axolotl, is a salamander that captivates scientists due to its exceptional characteristics (Bölük et al., 2022). Unlike other amphibians, axolotls exhibit neoteny, maintaining larval features while achieving sexual maturity (Sámano et al., 2021). However, their most remarkable attribute is the ability to regenerate nearly all their tissues and organs, setting them apart from other vertebrates. Despite descriptive and histological studies shedding light on tissue regeneration in *A. mexicanum*, the underlying molecular mechanisms remain enigmatic (S. V. Bryant et al., 2004; Knapp et al., 2013; Kragl et al., 2009). Comprehensive studies of the molecular mechanisms implicated in axolotl regeneration are of particular interest to the field of regenerative medicine, including potential implications for human therapies (Polikarpova et al., 2022; Samos et al., 2022). To gain a broader understanding of the genetic basis of axolotl tissue regeneration, we conducted a transcriptomic study on a group of *A. mexicanum* from Centro de Investigaciones Biológicas y Acuícolas de Cuemanco (CIBAC), a center dedicated to preserving the native axolotl population in Lake Xochimilco, Mexico City. Notably, these specimens were bred under conditions closely resembling their natural environment, reducing confounding factors that may be encountered in laboratory-raised counterparts (Reiß, 2022; Voss et al., 2015).

We began by examining samples from amputated limbs and the blastemas form in response to injury. Blastema formation is triggered by signals from surrounding tissues that induce the dedifferentiation of nearby cells into a proliferative and multipotent state. This organ is crucial for regeneration in axolotls, and because of that, is subject of a complex transcriptomic regulation (McCusker et al., 2015; Min & Whited, 2023). Employing next-generation RNA sequencing, we conducted differential expression analysis between blastemas and juvenile limbs, comparing our results to a previously published dataset to identify shared expression patterns. Additionally, we generated a predicted interactome and performed gene ontology annotation for the axolotl proteome to further describe the putative functions of the genes found.

Taken together, our findings suggest a global downregulation of genes associated with muscular tissue and anatomical development during limb regeneration, in line with cellular dedifferentiation that takes place in blastema (Gerber et al., 2018). However, a few genes related to development, such as *TBX4 (Khan et al., 2002)*, *BMP2 (Guimond et al., 2010)* and *KAZALD1* (D. M. Bryant et al., 2017), and wound healing, such as *SALL4 (Erickson et al., 2016)* were upregulated in blastemas. Furthermore, genes associated with the regulation of cell differentiation, such as *WNT5A* and *WNT5B*, exhibited increased expression. *WNT5A* and *WNT5B* have established roles in bone morphogenesis, hematopoiesis, and cartilage homeostasis (Mastelaro de Rezende et al., 2020; Suthon et al., 2021; Xuan et al., 2019). This suggests that, while tissue-specific genes are silenced during regeneration, a select few guide the development of the new limb.

In support of the aforementioned hypothesis, our gene coexpression network analysis reveals two modules that exhibit positive associations with blastemas samples. These modules contained several proteins involved in histone modification and DNA methylation, including *DNMT3A*, *HDAC2*, *SUV39H1*, *KDM3A* and the Polycomb group members *EED* and *EZH2*. Epigenetic regulation has been shown to be crucial for tissue regeneration in axolotls, pharmacological inhibition of DNA methyltransferases causes impairments in blastema formation (Aguilar & Gardiner, 2015) and histone deacetylase inhibitors inhibit regeneration (Voss et al., 2021). The Polycomb group is also known for its role in embryonic development and tissue differentiation, suggesting its likely importance in tissue regeneration for axolotls due to its evolutionary conservation (Whitcomb et al., 2007).

Intriguingly, we also observed upregulation of genes associated with extracellular matrix organization in the blastema compared to control tissue, while cell adhesion factors were predominantly downregulated. This may be linked to the extracellular matrix degradation (histolysis) that occurs at the amputation site, facilitating cell migration, differentiation, and blastema formation (Stocum, 2017). Among the main players in this tissue remodeling process are metalloproteases, with several, such as *MMP13*, *MMP19*, *ADAM8*, and *ADAMTS7*, found to be upregulated in the blastema (Santosh et al., 2011). Furthermore, genes involved in angiogenesis and wound healing, including tenascin (*TNC*), fibronectin (*FN1*), and thrombospondin-2 (*THBS2*), are also upregulated. Alongside the overall downregulation of myosin and actin proteins, these findings provide indicators of the extensive histolysis and angiogenesis process taking place in the tissue surrounding the blastema (Ritenour & Dickie, 2017; Whited et al., 2011).

The significance of the extracellular matrix becomes evident in the analysis of samples collected from aged axolotls that lacked regenerative capabilities. Notably, we detected the downregulation of several collagens and ribosomal proteins in aged limbs, indicative of impaired cell-cell interactions and extracellular matrix composition, commonly associated with aging (López-Otín et al., 2023; Selman & Pardo, 2021). Processes such as osteogenesis, proliferation and differentiation are influenced by the extracellular matrix stiffness (Du et al., 2016; Klein et al., 2009; Zhou et al., 2021), making the reduced collagen expression in aged organisms particularly intriguing. This leads us to hypothesize that extracellular matrix organization plays a central role in axolotl regeneration. Further investigation into the changes occurring in the extracellular matrix during blastema formation and limb regeneration in axolotls could provide valuable insights into the regulatory networks involved in this process.

A comparative analysis between aged limbs and blastema in juvenile specimens post-amputation led us to the identification of a set of genes that may be required for initiating the regeneration process. Among these, seven genes were pinpointed, including one putative lncRNA (*LOC112547415.222*) and six coding genes. A blast search against model animals demonstrated homologous counterparts of the six coding proteins in vertebrates, indicating their potential importance. Particularly, four regeneration-associated genes (*FSTL1*, *ADAMTS17*, *GPX7*, and *CTHRC1*) exhibited high expression levels in regenerating tissue but were underexpressed in aged axolotls. Structural and conservation analyses further highlighted that these genes encode conserved proteins in vertebrates, which together with structural predictions, allows us to infer their potential functions.

In this regard, ADAMTS17 is an extracellular metalloprotease involved in collagen processing, extracellular matrix degradation, cartilage cleavage, development, and angiogenesis. This protein has a conserved catalytic domain (Jones & Riley, 2005; Porter et al., 2005; Tang, 2001) which suggests it is catalytically active. Studies in *Adamts17* knockout mice showed that it is required for proper skeletogenesis and skeletal muscle development (Oichi et al., 2019). Mutations in ADAMTS17 are also linked to the Weill-Marchesani syndrome in humans, which affects connective tissue and leads to impaired vision, short stature, and musculoskeletal anomalies (Marzin et al., 2007; Yu et al., 2022). Additionally, ADAMTS17 mutations have been implicated in short height and glaucoma in dogs (Jeanes et al., 2019) and humans (van Duyvenvoorde et al., 2014). Nevertheless, the precise function of this protein remains enigmatic. Studies have observed that a catalytically inactive ADAMTS17 interferes with fibrillin-1 secretion, resulting in elastic fiber abnormalities and intracellular collagen accumulation in fibroblasts from patients with Weill-Marchesani syndrome (Karoulias et al., 2020). Another noteworthy protein involved in cell-matrix interactions is FSTL1, this is a secreted glycoprotein that participates in the regulation of the TGF-β, BMP and Wnt pathways (Hambrock et al., 2004; Horak et al., 2022). In axolotls, the identification of two domains, a Kasal and an EF-hand domain, aligns with the architecture described in other organisms (Li et al., 2019; Sylva et al., 2013). FSTL1 has been found to be critical for tracheal and central nervous system development in mice, with *Fstl1^-/-^*mice exhibiting cyanotic traits due to tracheal malformations (Geng et al., 2011; Yang et al., 2009). It is also involved in vascularization and vascular epithelial homeostasis maintenance (Jiang et al., 2020; Ouchi et al., 2008). Overall, the observed downregulation of ADAMTS17 and FSTL1 in aged axolotls compared to juvenile limbs further highlights the importance of the extracellular environment during limb regeneration.

Furthermore, CTHRC1, involved in the TGF-β pathway, functions as a mediator of osteoblast-osteoclast communication and plays a role in osteogenesis and bone remodeling (Kim et al., 2020). In axolotls, bone resorption mediated by osteoclast is important for adequate integration of regenerated bone (Riquelme-Guzmán et al., 2022). CTHRC1 also promotes angiogenesis by inhibiting collagen deposition and promoting cell migration (Myngbay et al., 2021). Knockout mice for this gene showed reduced bone density and arthritis (Jin et al., 2017; Takeshita et al., 2013), which also hints towards the importance of this protein for bone and cartilage development.

GPX7, another protein with high expression in blastema but reduced levels in aged limbs, is a peroxidase, and its catalytic domain has been found to be conserved in axolotls. GPX7 is vital for oxidative stress resistance (Bosello-Travain et al., 2013; D. Peng et al., 2012). Studies have linked low GPX7 levels to increased adipogenesis and fat accumulation in mice (Chang et al., 2013). Interestingly, a recent study observed that the bone marrow of aged axolotls has higher fat content than its younger counterpart (Polikarpova et al., 2022), which could align with the reduced GPX7 expression in aged axolotl limbs. Moreover, tissue regeneration in axolotls requires the production of reactive oxygen species (Al Haj Baddar et al., 2019), suggesting that enzymes involved in the regulation of oxidative stress are important for this process.

Lastly, we also identified two nicotinamide *N*-methyltransferases (NNMT) proteins with high expression in aged axolotls but with decreased expression in blastema. While the sequence of both proteins showed homology to other vertebrate NNMTs, we noticed that only half of the catalytic domain of the protein is present in our axolotl proteins. NNMTs enzymes belong to the group of SAM-dependent methyltransferases (Struck et al., 2012), and they have been implicated in several epigenetic processes since SAM serves as the methyl donor used by DNA and histone methyltransferases. Consequently, high levels of NNMT cause a decrease in SAM availability and in consequence, leading to the inhibition of other methyltransferases (Eckert et al., 2019; Sperber et al., 2015). Although the functional status of the axolotl NNMT proteins, as identified by us, is not clear, it is evident that these transcripts play a central role in aged axolotls, and further investigation is needed to unravel their mechanisms of action. These two transcripts were identified in the yellow co-expression module, displaying a strong negative correlation with blastema and enrichment in genes associated with metabolic processes, thereby accentuating their central role in regeneration and aged limbs.

In summary, with this work we present a broad description of the transcriptional changes that take place during tissue regeneration in axolotls, and with the aid of aged axolotl samples we identified a set of genes that may be crucial for the regenerative capabilities of *A. mexicanum*. Through conservation analysis, we suggest that these genes have important roles in development, bone morphogenesis, and extracellular matrix organization. Our results also showed that the proteins coded by these genes can be found in other vertebrates, thus, could be used as a starting point for the study of regeneration in other animals. Our findings highlight the importance of understanding the transcriptomic landscape of A. mexicanum and provide valuable insights into the molecular mechanisms underlying tissue regeneration and the effects of aging on regenerative capacity.

## Methods

### Sample collection

All biological samples were obtained from a captive population of Mexican axolotls (*Ambystoma mexicanum)*, residing at the Unidad de Manejo Ambiental of the Centro de Investigaciones Biológicas y Acuícolas de Cuemanco (UMA-CIBAC), which is part of the Universidad Autónoma Metropolitana, Xochimilco Campus (UAM-Xochimilco) in Mexico City. The protocol followed the guidelines established by UMA-CIBAC and UAM-Xochimilco. All the procedures were approved and supervised by the veterinarians from UMA-CIBAC, according to the norms and regulations set by the Mexican Ministry of Environment and Natural Resources. Sedation of the axolotls was accomplished through immersion in a tank containing benzocaine at a concentration of 50 mg/L prior to the amputation process. Biological samples were meticulously collected, using a stereoscopic microscope, from the inferior limbs of five juvenile axolotls, aged 8 months, as well as from two aged axolotls, aged 8 years. Specifically, 10 days post-amputation, blastema tissues were acquired from juvenile axolotls, whereas aged axolotls did not display any development of blastema tissue. No animals were sacrificed for the purpose of this study, and the axolotls were safely reintroduced to their habitat at UMA-CIBAC following the final sample collection.

### RNA extraction and sequencing

The collected tissue was preserved in RNA*later* (Invitrogen), at 4°C for a maximum of 24 hrs prior to processing. The RNA was extracted according to the protocol established by Peña-Llopis et al (Peña-Llopis & Brugarolas, 2013). RNA quality and concentration were assessed using the High Sensitivity RNA Tapestation (Agilent Technologies). Ribosomal RNA depletion was performed with the RiboZero Gold kit (Illumina), and libraries were constructed using the SmarterStranded V2 kit (Takara Bio). Paired-end (PE) sequencing was carried out on Illumina with a read length configuration of 150 and 20 million reads per sample.

### Transcriptomic analysis

In addition to the experimentally obtained samples, the data from two other studies was also analyzed. The accession numbers for the samples used are displayed in **Table 3**.

**Table 3.**
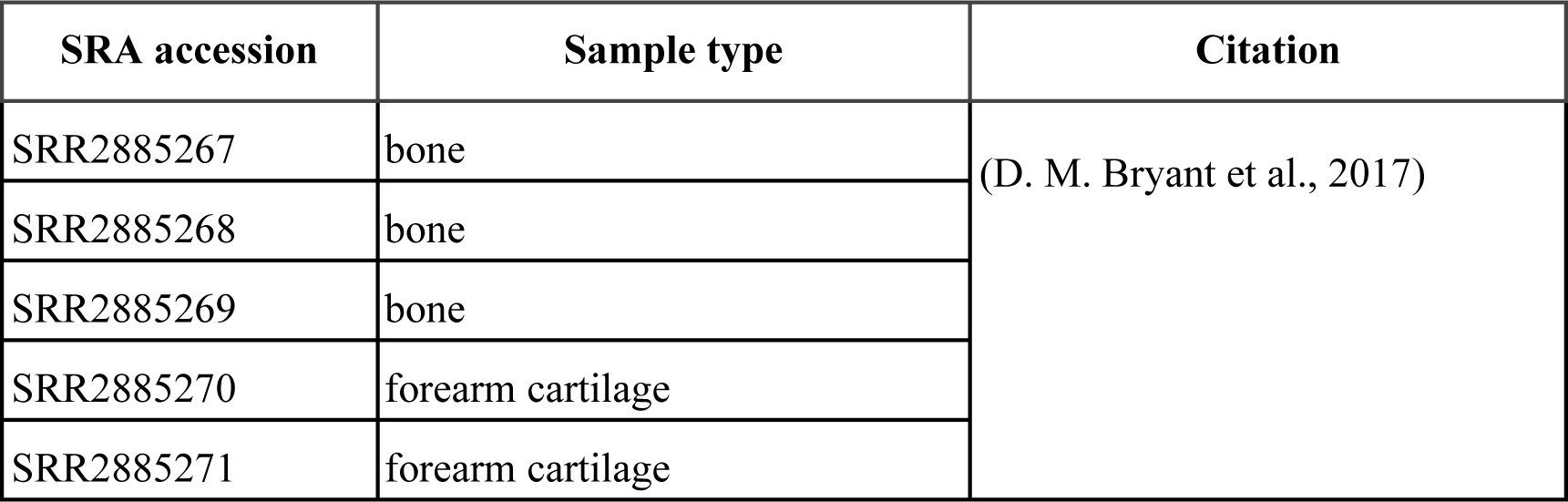
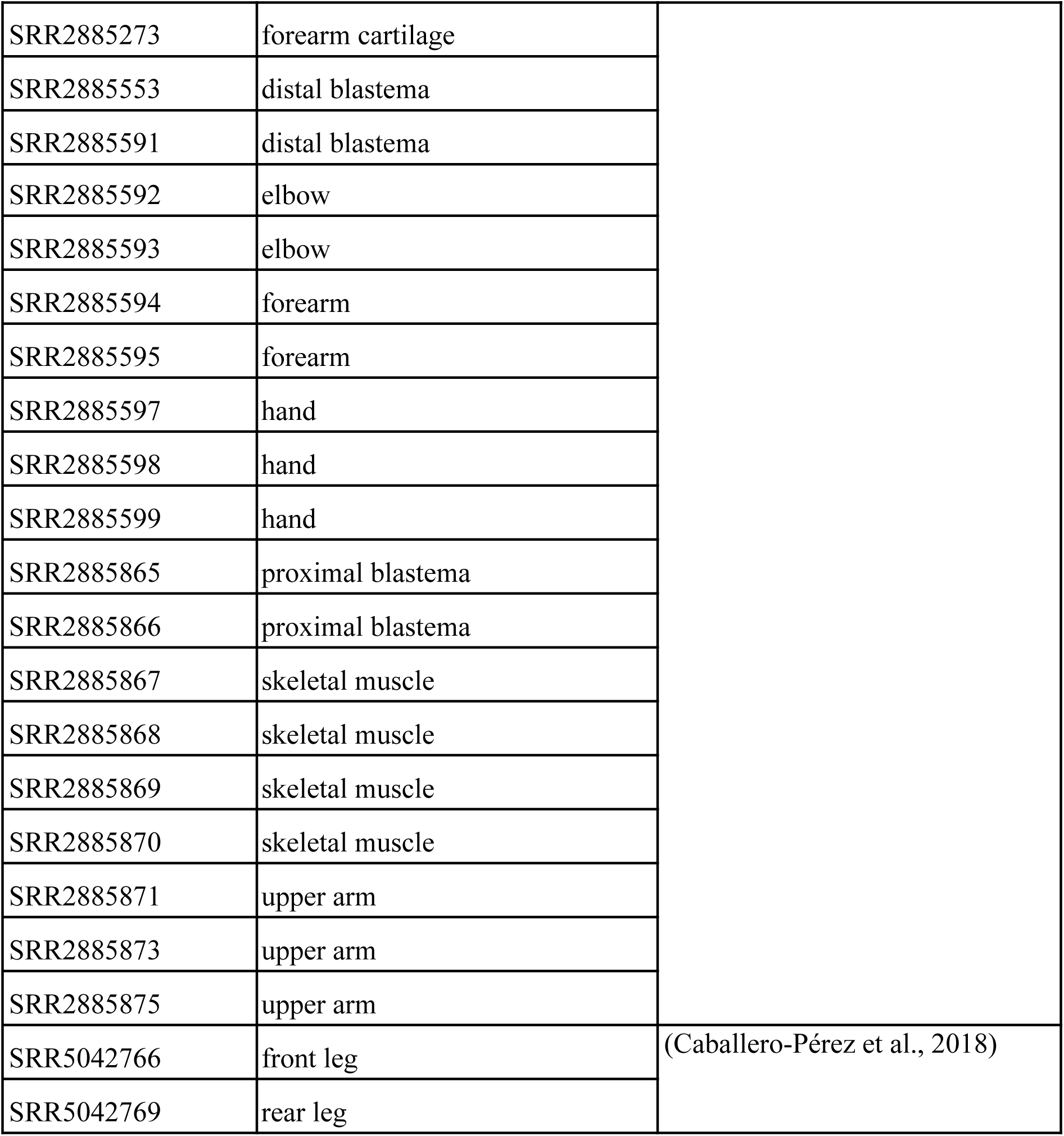
Previously published samples analyzed.

Quality assessment of the sequencing data was performed using FastQC (Andrews, 2010) Subsequently, raw counts were aligned to the reference genome AmexG_v6.0-DD (Schloissnig et al., 2021) using STAR v.2.7.9a (Dobin et al., 2013). The resulting transcript-aligned read counts were quantified using featureCounts for R, and the AmexT_v47 transcriptome, accessible at https://www.axolotl-omics.org/assemblies, was used for this purpose. DESeq2 v.1.32.0 (Love et al., 2014) was employed for the differential expression analysis. The tissue obtained after amputation of the juvenile axolotl’s limb was used as a control. Two separate comparisons were carried out: aged axolotl’s limb (8 years old) versus control, and 10-day blastema from juvenile axolotls versus control. Genes were designated as differentially expressed (DEG) if they exhibited |log2(FoldChange)| > 1 and padj < 0.05. For the Bryant et al. dataset, differential expression analysis was conducted for proximal blastema using upper arm tissue as the control, and for distal blastema using hand tissue as the control.

### Gene annotation and protein-protein interaction networks

All genes were named according to the “gene_name” field in the AmexT_v47 transcriptome file, which can be accessed from https://www.axolotl-omics.org/assemblies. In cases where the “gene_name” field was either empty or equal to “N/A,” genes were assigned the same name as their gene_id. Priority was given to nomenclature with the [hs] identifier over any other annotation. To distinguish genes with the same name, a numerical identifier was appended, e.g., KAZAL.3 indicated the presence of four other genes with the name KAZAL in the dataset. The predicted Gene Ontology (GO) function and protein-protein interactions (PPI) for each gene were determined using STRING (Szklarczyk et al., 2023). Only one open read frame per gene was used as input for STRING; if a gene had more than one transcript, the .1 isoform was selected. The GO term enrichment analysis was performed using gProfiler2 v.0.2.1 for R (Kolberg et al., 2020) with the correction method g:SCS and an adjusted p-value significance threshold of 0.05. Redundant terms were clustered using rrvgo v.1.4.4 for R (Sayols, 2023). The PPI visualizations were built with KeyPathwayMineR (Mechteridis et al., 2022) All network visualizations were created with Cytoscape (Bindea et al., 2009).

### Co-expression analysis

Our datasets, as well as the ones from Bryant and Caballero-Pérez, were used as input for WGCNA(Langfelder & Horvath, 2008). The raw counts were normalized using variance-stabilizing transformation (vst) from the DESeq2 package. Batch effect correction was performed with Combat, from the sva v3.40.0 package for R(Johnson et al., 2007). The blockwiseModules function was employed to identify co-expression modules within the data. The function was run with a power of 7 and a minimum module size of 30 genes, the TOMType was set to “unsigned”. The resulting co-expression modules were named with random colors. Modules were considered as significant if they had a p-value < 0.05 and |Module-Trait Correlation| > 0.5. The GO enrichment analysis for the genes contained in each module was performed with gProfiler2 v.0.2.1 for R(Kolberg et al., 2020) and the protein-protein interaction network was built with KeyPathwayMineR(Mechteridis et al., 2022) using only the genes with Module Membership > 0.7. The visualizations were created with Cytoscape and BioRender.com.

### Homology and structural biology analysis

The amino acid sequences for each transcript were obtained from the annotation file (.gtf) for the AmexT_v47 transcriptome. If more than one transcript was available, only the .1 isoform was modeled. The amino acid sequence of each protein was used as a query for BLASTP 2.12.0 (Altschul et al., 1997; Schäffer et al., 2001) against the UniProtKB and Swiss-Prot database (UniProt Consortium, 2023). Only the sequence with the highest identity percentage in each organism was conserved for the identity plot, and the organisms displayed were selected based on the animal model organisms described in a comprehensive review about model organisms (Hedges, 2002). AlphaFold2 (Jumper et al., 2021) was used to predict the structures of the proteins, templates were used for all predictions, and the ConSurf (Yariv et al., 2023) web server was used to color the residues according to their conservation. The functional domain annotation was performed using the Conserved Domain Database (CDD) (Marchler-Bauer et al., 2015) and the NCBI Conserved Domain tool (Marchler-Bauer & Bryant, 2004). The Protein visualizations were created with the PyMOL Molecular Graphics System v.4.60.

## Data availability

The GO term information and PPI network built and used in this work can be explored in STRING using the organism identifier STRG0034MNQ or the link https://version-11-5.string-db.org/organism/STRG0034MNQ. Additionally, the gene annotation file, ppi predicted network, an AnnotationDbi OrgDb package and a Txdb object with the gene names and associated GO terms can be accessed at the https://github.com/aylindmm/A.mexicanum GitHub repository. The raw sequencing data was deposited in NCBI’s Gene Expression Omnibus under the accession id <u>GSE237864</u>.

## Ethics statement

All studies conducted with *Ambystoma mexicanum* were approved under the ethical considerations authorized by the ethics and research committee (Reference Number: CEI.2023.007) of the Division of Biological and Health Sciences at the Universidad Autónoma Metropolitana, Xochimilco Campus, according to the current legislation and the guidelines established by the Mexican Ministry of Environment and Natural Resources.

## Acknowledgements

We would like to thank M. Sc. Karla Torres-Arciga and Michel Montalvo Casimiro for their technical support in processing the RNA samples of *Ambystoma mexicanum*.

This work was funded by the Consejo Nacional de Ciencia y Tecnología (CONACyT) Fondo CB-SEP-CONACyT 284748 and PRODEP (511/2023-3066-1599) to E.S-R. All the computational analyses were performed on the esrs-epigenetics server from Universidad Autónoma Metropolitana (UAM)-Cuajimalpa funded by CONACyT through the Apoyo para proyectos de investigación científica, desarrollo tecnológico e innovación en salud ante la contingencia por COVID-19, grant number 00312021. C.S. and E.S-R were supported by the Departamento de Ciencias Naturales (DCN) de la División de Ciencias Naturales e Ingeniería (DCNI) de la UAM-Cuajimalpa by Divisional Project Number 47301025. A.D.M.M. is a doctoral student from “Programa de maestría y doctorado en Ciencias Bioquímicas”, Universidad Nacional Autónoma de México (UNAM) and received a PhD fellowship funding from CONACyT (CVU 894530) and a Research Grant for Bi-nationally Supervised Doctoral Degrees/Cotutelle from Deutscher Akademischer Austauschdienst (DAAD, Personal reference: 91833882). The work of J.B. and M.T. was developed as part of the ASPIRE and SyMBoD projects and is funded by the German Federal Ministry of Education and Research (BMBF) under grant numbers 031L0287B and 01ZX1910D. M.T was partly funded by the Leibniz ScienceCampus InterACt (from BWFGB Hamburg and the Leibniz Association).

## Author contributions

Conceptualization, C.S. and E.S.R.; Methodology, C.S., E.S.R., J.A.O.C and R.G.B.; Investigation, C.S., E.S.R., J.A.O.C, and A.D.M.M.; Formal Analysis, A.D.M.M.; Data Curation, A.D.M.M; Writing – Original Draft, A.D.M.M.; Writing – Review & Editing, C.S. E.S.R, J.B. and M.T.; Funding Acquisition, E.S.R, J.B. and M.T.; Resources, E.S.R., R.G.B., and J.A.O.C; Supervision, C.S., J.B., M.T and E.S.R.

## References

Aguilar, C., & Gardiner, D. M. (2015). DNA Methylation Dynamics Regulate the Formation of a Regenerative Wound Epithelium during Axolotl Limb Regeneration. PloS One, 10(8), e0134791.

Al Haj Baddar, N. W., Chithrala, A., & Voss, S. R. (2019). Amputation-induced reactive oxygen species signaling is required for axolotl tail regeneration. Developmental Dynamics: An Official Publication of the American Association of Anatomists, 248(2), 189–196.

Altschul, S. F., Madden, T. L., Schäffer, A. A., Zhang, J., Zhang, Z., Miller, W., & Lipman, D. J. (1997). Gapped BLAST and PSI-BLAST: a new generation of protein database search programs. Nucleic Acids Research, 25(17), 3389–3402.

Andrews, S. (2010). FastQC: A Quality Control Tool for High Throughput Sequence Data. http://www.bioinformatics.babraham.ac.uk/projects/fastqc/

Bindea, G., Mlecnik, B., Hackl, H., Charoentong, P., Tosolini, M., Kirilovsky, A., Fridman, W.-H., Pagès, F., Trajanoski, Z., & Galon, J. (2009). ClueGO: a Cytoscape plug-in to decipher functionally grouped gene ontology and pathway annotation networks. Bioinformatics, 25(8), 1091–1093.

Bölük, A., Yavuz, M., & Demircan, T. (2022). Axolotl: A resourceful vertebrate model for regeneration and beyond. Developmental Dynamics: An Official Publication of the American Association of Anatomists, 251(12), 1914–1933.

Bosello-Travain, V., Conrad, M., Cozza, G., Negro, A., Quartesan, S., Rossetto, M., Roveri, A., Toppo, S., Ursini, F., Zaccarin, M., & Maiorino, M. (2013). Protein disulfide isomerase and glutathione are alternative substrates in the one Cys catalytic cycle of glutathione peroxidase 7. Biochimica et Biophysica Acta, 1830(6), 3846–3857.

Bryant, D. M., Johnson, K., DiTommaso, T., Tickle, T., Couger, M. B., Payzin-Dogru, D., Lee, T. J., Leigh, N. D., Kuo, T.-H., Davis, F. G., Bateman, J., Bryant, S., Guzikowski, A. R., Tsai, S. L., Coyne, S., Ye, W. W., Freeman, R. M., Jr, Peshkin, L., Tabin, C. J., … Whited, J. L. (2017). A Tissue-Mapped Axolotl De Novo Transcriptome Enables Identification of Limb Regeneration Factors. Cell Reports, 18(3), 762–776.

Bryant, S. V., Endo, T., & Gardiner, D. M. (2004). Vertebrate limb regeneration and the origin of limb stem cells. The International Journal of Developmental Biology, 46(7), 887–896.

Caballero-Pérez, J., Espinal-Centeno, A., Falcon, F., García-Ortega, L. F., Curiel-Quesada, E., Cruz-Hernández, A., Bako, L., Chen, X., Martínez, O., Alberto Arteaga-Vázquez, M., Herrera-Estrella, L., & Cruz-Ramírez, A. (2018). Transcriptional landscapes of Axolotl (Ambystoma mexicanum). Developmental Biology, 433(2), 227–239.

Chang, Y.-C., Yu, Y.-H., Shew, J.-Y., Lee, W.-J., Hwang, J.-J., Chen, Y.-H., Chen, Y.-R., Wei, P.-C., Chuang, L.-M., & Lee, W.-H. (2013). Deficiency of NPGPx, an oxidative stress sensor, leads to obesity in mice and human. EMBO Molecular Medicine, 5(8), 1165–1179.

Darnet, S., Dragalzew, A. C., Amaral, D. B., Sousa, J. F., Thompson, A. W., Cass, A. N., Lorena, J., Pires, E. S., Costa, C. M., Sousa, M. P., Fröbisch, N. B., Oliveira, G., Schneider, P. N., Davis, M. C., Braasch, I., & Schneider, I. (2019). Deep evolutionary origin of limb and fin regeneration. Proceedings of the National Academy of Sciences of the United States of America, 116(30), 15106–15115.

Demircan, T., İlhan, A. E., Aytürk, N., Yıldırım, B., Öztürk, G., & Keskin, İ. (2016). A histological atlas of the tissues and organs of neotenic and metamorphosed axolotl. Acta Histochemica, 118(7), 746–759.

Dobin, A., Davis, C. A., Schlesinger, F., Drenkow, J., Zaleski, C., Jha, S., Batut, P., Chaisson, M., & Gingeras, T. R. (2013). STAR: ultrafast universal RNA-seq aligner. Bioinformatics, 29(1), 15–21.

Du, J., Zu, Y., Li, J., Du, S., Xu, Y., Zhang, L., Jiang, L., Wang, Z., Chien, S., & Yang, C. (2016). Extracellular matrix stiffness dictates Wnt expression through integrin pathway. Scientific Reports, 6, 20395.

Eckert, M. A., Coscia, F., Chryplewicz, A., Chang, J. W., Hernandez, K. M., Pan, S., Tienda, S. M., Nahotko, D. A., Li, G., Blaženović, I., Lastra, R. R., Curtis, M., Yamada, S. D., Perets, R., McGregor, S. M., Andrade, J., Fiehn, O., Moellering, R. E., Mann, M., & Lengyel, E. (2019). Proteomics reveals NNMT as a master metabolic regulator of cancer-associated fibroblasts. Nature, 569(7758), 723–728.

Erickson, J. R., Gearhart, M. D., Honson, D. D., Reid, T. A., Gardner, M. K., Moriarity, B. S., & Echeverri, K. (2016). A novel role for SALL4 during scar-free wound healing in axolotl. NPJ Regenerative Medicine, 1, 16016 – .

Farkas, J. E., & Monaghan, J. R. (2015). Housing and maintenance of Ambystoma mexicanum, the Mexican axolotl. Methods in Molecular Biology, 1290, 27–46.

Geng, Y., Dong, Y., Yu, M., Zhang, L., Yan, X., Sun, J., Qiao, L., Geng, H., Nakajima, M., Furuichi, T., Ikegawa, S., Gao, X., Chen, Y.-G., Jiang, D., & Ning, W. (2011). Follistatin-like 1 (Fstl1) is a bone morphogenetic protein (BMP) 4 signaling antagonist in controlling mouse lung development. Proceedings of the National Academy of Sciences of the United States of America, 108(17), 7058–7063.

Gerber, T., Murawala, P., Knapp, D., Masselink, W., Schuez, M., Hermann, S., Gac-Santel, M., Nowoshilow, S., Kageyama, J., Khattak, S., Currie, J. D., Camp, J. G., Tanaka, E. M., & Treutlein, B. (2018). Single-cell analysis uncovers convergence of cell identities during axolotl limb regeneration. Science, 362(6413). 10.1126/science.aaq0681

Guimond, J.-C., Lévesque, M., Michaud, P.-L., Berdugo, J., Finnson, K., Philip, A., & Roy, S. (2010). BMP-2 functions independently of SHH signaling and triggers cell condensation and apoptosis in regenerating axolotl limbs. BMC Developmental Biology, 10, 15.

Haas, B. J., & Whited, J. L. (2017). Advances in Decoding Axolotl Limb Regeneration. Trends in Genetics: TIG, 33(8), 553–565.

Hambrock, H. O., Kaufmann, B., Müller, S., Hanisch, F.-G., Nose, K., Paulsson, M., Maurer, P., & Hartmann, U. (2004). Structural Characterization of TSC-36/Flik. Journal of Biological Chemistry, 279(12), 11727–11735.

Hedges, S. B. (2002). The origin and evolution of model organisms. Nature Reviews. Genetics, 3(11), 838–849.

Horak, M., Fairweather, D., Kokkonen, P., Bednar, D., & Bienertova-Vasku, J. (2022). Follistatin-like 1 and its paralogs in heart development and cardiovascular disease. Heart Failure Reviews, 27(6), 2251–2265.

Jeanes, E. C., Oliver, J. A. C., Ricketts, S. L., Gould, D. J., & Mellersh, C. S. (2019). Glaucoma-causing mutations are also reproducibly associated with height in two domestic dog breeds: selection for short stature may have contributed to increased prevalence of glaucoma. Canine Genetics and Epidemiology, 6, 5.

Jiang, H., Zhang, L., Liu, X., Sun, W., Kato, K., Chen, C., Li, X., Li, T., Sun, Z., Han, W., Zhang, F., Xiao, Q., Yang, Z., Hu, J., Qin, Z., Adams, R. H., Gao, X., & He, Y. (2020). Angiocrine FSTL1 (Follistatin-Like Protein 1) Insufficiency Leads to Atrial and Venous Wall Fibrosis via SMAD3 Activation. *Arteriosclerosis*, Thrombosis, and Vascular Biology, 40(4), 958–972.

Jin, Y.-R., Stohn, J. P., Wang, Q., Nagano, K., Baron, R., Bouxsein, M. L., Rosen, C. J., Adarichev, V. A., & Lindner, V. (2017). Inhibition of osteoclast differentiation and collagen antibody-induced arthritis by CTHRC1. Bone, 97, 153–167.

Johnson, W. E., Li, C., & Rabinovic, A. (2007). Adjusting batch effects in microarray expression data using empirical Bayes methods. Biostatistics, 8(1), 118–127.

Jones, G. C., & Riley, G. P. (2005). ADAMTS proteinases: a multi-domain, multi-functional family with roles in extracellular matrix turnover and arthritis. Arthritis Research & Therapy, 7(4), 160–169.

Joven, A., Elewa, A., & Simon, A. (2019). Model systems for regeneration: salamanders.Development, 146(14). 10.1242/dev.167700

Jumper, J., Evans, R., Pritzel, A., Green, T., Figurnov, M., Ronneberger, O., Tunyasuvunakool, K., Bates, R., Žídek, A., Potapenko, A., Bridgland, A., Meyer, C., Kohl, S. A. A., Ballard, A. J., Cowie, A., Romera-Paredes, B., Nikolov, S., Jain, R., Adler, J., … Hassabis, D. (2021). Highly accurate protein structure prediction with AlphaFold. Nature, 596(7873), 583–589.

Karoulias, S. Z., Beyens, A., Balic, Z., Symoens, S., Vandersteen, A., Rideout, A. L., Dickinson, J., Callewaert, B., & Hubmacher, D. (2020). A novel ADAMTS17 variant that causes Weill-Marchesani syndrome 4 alters fibrillin-1 and collagen type I deposition in the extracellular matrix. Matrix Biology: Journal of the International Society for Matrix Biology, 88, 1–18.

Kavanagh, K. L., Johansson, C., Papagrigoriou, E., Kochan, G., Umeano, C., Gileadi, O., von Delft, F., Weigelt, J., Arrowsmith C. H., Sundstrom, M., Edwards, A., Oppermann, U., & Structural Genomics Consortium (SGC). (2007). PDB: 2P31 [Data set]. In Crystal structure of human glutathione peroxidase 7. 10.2210/pdb2p31/pdb

Keinath, M. C., Timoshevskiy, V. A., Timoshevskaya, N. Y., Tsonis, P. A., Voss, S. R., & Smith, J. J. (2015). Initial characterization of the large genome of the salamander Ambystoma mexicanum using shotgun and laser capture chromosome sequencing. Scientific Reports, 5, 16413.

Khan, P., Linkhart, B., & Simon, H.-G. (2002). Different regulation of T-box genes Tbx4 and Tbx5 during limb development and limb regeneration. Developmental Biology, 250(2), 383–392.

Kim, J.-M., Lin, C., Stavre, Z., Greenblatt, M. B., & Shim, J.-H. (2020). Osteoblast-Osteoclast Communication and Bone Homeostasis. Cells, 9(9). 10.3390/cells9092073

Klein, E. A., Yin, L., Kothapalli, D., Castagnino, P., Byfield, F. J., Xu, T., Levental, I., Hawthorne, E., Janmey, P. A., & Assoian, R. K. (2009). Cell-cycle control by physiological matrix elasticity and in vivo tissue stiffening. Current Biology: CB, 19(18), 1511–1518.

Knapp, D., Schulz, H., Rascon, C. A., Volkmer, M., Scholz, J., Nacu, E., Le, M., Novozhilov, S., Tazaki, A., Protze, S., Jacob, T., Hubner, N., Habermann, B., & Tanaka, E. M. (2013). Comparative transcriptional profiling of the axolotl limb identifies a tripartite regeneration-specific gene program. PloS One, 8(5), e61352.

Kolberg, L., Raudvere, U., Kuzmin, I., & Vilo, J. (2020). gprofiler2--an R package for gene list functional enrichment analysis and namespace conversion toolset g: Profiler. F1000Research, 9(ELIXIR-709). https://www.ncbi.nlm.nih.gov/pmc/articles/PMC7859841.2/

Kragl, M., Knapp, D., Nacu, E., Khattak, S., Maden, M., Epperlein, H. H., & Tanaka, E. M. (2009). Cells keep a memory of their tissue origin during axolotl limb regeneration. Nature, 460(7251), 60–65.

Langfelder, P., & Horvath, S. (2008). WGCNA: an R package for weighted correlation network analysis. BMC Bioinformatics, 9(1), 1–13.

Leclère, L., Nir, T. S., Bazarsky, M., Braitbard, M., Schneidman-Duhovny, D., & Gat, U. (2020). Dynamic Evolution of the Cthrc1 Genes, a Newly Defined Collagen-Like Family. Genome Biology and Evolution, 12(2), 3957–3970.

Lin, T. Y., Gerber, T., Taniguchi-Sugiura, Y., Murawala, P., Hermann, S., Grosser, L., Shibata, E., Treutlein, B., & & Tanaka, E. M. (2021). Fibroblast dedifferentiation as a determinant of successful regeneration. Developmental Cell, 56(10), 1541–1551.e6.

Li, X., Li, L., Chang, Y., Ning, W., & Liu, X. (2019). Structural and functional study of FK domain of Fstl1. Protein Science: A Publication of the Protein Society, 28(10), 1819–1829.

López-Otín, C., Blasco, M. A., Partridge, L., Serrano, M., & Kroemer, G. (2023). Hallmarks of aging: An expanding universe. Cell, 186(2), 243–278.

Love, M. I., Huber, W., & Anders, S. (2014). Moderated estimation of fold change and dispersion for RNA-seq data with DESeq2. Genome Biology, 15(12), 550.

Makanae, A., Mitogawa, K., & Satoh, A. (2014). Co-operative Bmp- and Fgf-signaling inputs convert skin wound healing to limb formation in urodele amphibians. Developmental Biology, 396(1), 57–66.

Marchler-Bauer, A., & Bryant, S. H. (2004). CD-Search: protein domain annotations on the fly. Nucleic Acids Research, 32(Web Server issue), W327–W331.

Marchler-Bauer, A., Derbyshire, M. K., Gonzales, N. R., Lu, S., Chitsaz, F., Geer, L. Y., Geer, R. C., He, J., Gwadz, M., Hurwitz, D. I., Lanczycki, C. J., Lu, F., Marchler, G. H., Song, J. S., Thanki, N., Wang, Z., Yamashita, R. A., Zhang, D., Zheng, C., & Bryant, S. H. (2015). CDD: NCBI’s conserved domain database. Nucleic Acids Research, 43(Database issue), D222–D226.

Marzin, P., Cormier-Daire, V., & Tsilou, E. (2007). Weill-Marchesani Syndrome. In M. P. Adam, G. M. Mirzaa, R. A. Pagon, S. E. Wallace, L. J. H. Bean, K. W. Gripp, & A. Amemiya (Eds.), GeneReviews. University of Washington, Seattle.

Masselink, W., & Tanaka, E. M. (2021). Toward whole tissue imaging of axolotl regeneration. Developmental Dynamics: An Official Publication of the American Association of Anatomists, 250(6), 800–806.

Mastelaro de Rezende, M., Zenker Justo, G., Julian Paredes-Gamero, E., & Gosens, R. (2020). Wnt-5A/B Signaling in Hematopoiesis throughout Life. Cells, 9(8). 10.3390/cells9081801

McCusker, C., Bryant, S. V., & Gardiner, D. M. (2015). The axolotl limb blastema: cellular and molecular mechanisms driving blastema formation and limb regeneration in tetrapods. *Regeneration (Oxford*, England), 2(2), 54–71.

McCusker, C., & Gardiner, D. M. (2011). The axolotl model for regeneration and aging research: a mini-review. Gerontology, 57(6), 565–571.

McHedlishvili, L., Mazurov, V., Grassme, K. S., Goehler, K., Robl, B., Tazaki, A., Roensch, K., Duemmler, A., & Tanaka, E. M. (2012). Reconstitution of the central and peripheral nervous system during salamander tail regeneration. Proceedings of the National Academy of Sciences of the United States of America, 109(34), E2258–E2266.

Mechteridis, K., Lauber, M., Baumbach, J., & List, M. (2022). KeyPathwayMineR: De Novo Pathway Enrichment in the R Ecosystem. Frontiers in Genetics, 12. 10.3389/fgene.2021.812853

Min, S., & Whited, J. L. (2023). Limb blastema formation: How much do we know at a genetic and epigenetic level? The Journal of Biological Chemistry, 299(2), 102858.

Monaghan, J. R., Athippozhy, A., Seifert, A. W., Putta, S., Stromberg, A. J., Maden, M., Gardiner, D. M., & Voss, S. R. (2012). Gene expression patterns specific to the regenerating limb of the Mexican axolotl. Biology Open, 1(10), 937–948.

Mosyak, L., Georgiadis, K., Shane, T., Svenson, K., Hebert, T., McDonagh, T., Mackie, S., Olland, S., Lin, L., Zhong, X., Kriz, R., Reifenberg, E. L., Collins-Racie, L. A., Corcoran, C., Freeman, B., Zollner, R., Marvell, T., Vera, M., Sum, P.-E., … Somers, W. (2008). Crystal structures of the two major aggrecan degrading enzymes, ADAMTS4 and ADAMTS5. Protein Science: A Publication of the Protein Society, 17(1), 16–21.

Myngbay, A., Manarbek, L., Ludbrook, S., & Kunz, J. (2021). The Role of Collagen Triple Helix Repeat-Containing 1 Protein (CTHRC1) in Rheumatoid Arthritis. International Journal of Molecular Sciences, 22(5). 10.3390/ijms22052426

Oichi, T., Taniguchi, Y., Soma, K., Oshima, Y., Yano, F., Mori, Y., Chijimatsu, R., Kim-Kaneyama, J.-R., Tanaka, S., & Saito, T. (2019). Adamts17 is involved in skeletogenesis through modulation of BMP-Smad1/5/8 pathway. Cellular and Molecular Life Sciences: CMLS, 76(23), 4795–4809.

Ouchi, N., Oshima, Y., Ohashi, K., Higuchi, A., Ikegami, C., Izumiya, Y., & Walsh, K. (2008). Follistatin-like 1, a secreted muscle protein, promotes endothelial cell function and revascularization in ischemic tissue through a nitric-oxide synthase-dependent mechanism. The Journal of Biological Chemistry, 283(47), 32802–32811.

Peña-Llopis, S., & Brugarolas, J. (2013). Simultaneous isolation of high-quality DNA, RNA, miRNA and proteins from tissues for genomic applications. Nature Protocols, 8(11), 2240–2255.

Peng, D., Belkhiri, A., Hu, T., Chaturvedi, R., Asim, M., Wilson, K. T., Zaika, A., & El-Rifai, W. (2012). Glutathione peroxidase 7 protects against oxidative DNA damage in oesophageal cells. Gut, 61(9), 1250–1260.

Peng, Y., Sartini, D., Pozzi, V., Wilk, D., Emanuelli, M., & Yee, V. C. (2011). Structural basis of substrate recognition in human nicotinamide N-methyltransferase. Biochemistry, 50(36), 7800–7808.

Polikarpova, A., Ellinghaus, A., Schmidt-Bleek, O., Grosser, L., Bucher, C. H., Duda, G. N., Tanaka, E. M., & Schmidt-Bleek, K. (2022). The specialist in regeneration-the Axolotl-a suitable model to study bone healing? NPJ Regenerative Medicine, 7(1), 35.

Porter, S., Clark, I. M., Kevorkian, L., & Edwards, D. R. (2005). The ADAMTS metalloproteinases. Biochemical Journal, 386(1), 15–27.

Reiß, C. (2022). Cut and Paste: The Mexican Axolotl, Experimental Practices and the Long History of Regeneration Research in Amphibians, 1864-Present. Frontiers in Cell and Developmental Biology, 10, 786533.

Riquelme-Guzmán, C., Tsai, S. L., Carreon Paz, K., Nguyen, C., Oriola, D., Schuez, M., Brugués, J., Currie, J. D., & Sandoval-Guzmán, T. (2022). Osteoclast-mediated resorption primes the skeleton for successful integration during axolotl limb regeneration. eLife, 11. 10.7554/eLife.79966

Ritenour, A. M., & Dickie, R. (2017). Inhibition of Vascular Endothelial Growth Factor Receptor Decreases Regenerative Angiogenesis in Axolotls. Anatomical Record, 300(12), 2273–2280.

Sámano, C., González-Barrios, R., Castro-Azpíroz, M., Torres-García, D., Ocampo-Cervantes, J. A., Otero-Negrete, J., & Soto-Reyes, E. (2021). Genomics and epigenomics of axolotl regeneration. The International Journal of Developmental Biology, 65(7-8-9), 465–474.

Samos, A., McGaughey, V., Rieger, S., & Lisse, T. S. (2022). Reawakening GDNF’s regenerative past in mice and humans. Regenerative Therapy, 20, 78–85.

Sandoval-Guzmán, T., Wang, H., Khattak, S., Schuez, M., Roensch, K., Nacu, E., Azaki, T., Joven, A., Tanaka, E. M., & Simon, A. (2014). Fundamental Differences in Dedifferentiation and Stem Cell Recruitment during Skeletal Muscle Regeneration in Two Salamander Species. Cell Stem Cell, 14(2), 174–187.

Santosh, N., Windsor, L. J., Mahmoudi, B. S., Li, B., Zhang, W., Chernoff, E. A., Rao, N., Stocum, D. L., & Song, F. (2011). Matrix metalloproteinase expression during blastema formation in regeneration-competent versus regeneration-deficient amphibian limbs. Developmental Dynamics: An Official Publication of the American Association of Anatomists, 240(5), 1127–1141.

Sayols, S. (2023). rrvgo: a Bioconductor package for interpreting lists of Gene Ontology terms. microPublication Biology, 2023. 10.17912/micropub.biology.000811

Schäffer, A. A., Aravind, L., Madden, T. L., Shavirin, S., Spouge, J. L., Wolf, Y. I., Koonin, E. V., & Altschul, S. F. (2001). Improving the accuracy of PSI-BLAST protein database searches with composition-based statistics and other refinements. Nucleic Acids Research, 29(14), 2994–3005.

Schloissnig, S., Kawaguchi, A., Nowoshilow, S., Falcon, F., Otsuki, L., Tardivo, P., Timoshevskaya, N., Keinath, M. C., Smith, J. J., Voss, S. R., & Tanaka, E. M. (2021). The giant axolotl genome uncovers the evolution, scaling, and transcriptional control of complex gene loci. Proceedings of the National Academy of Sciences of the United States of America, 118(15). 10.1073/pnas.2017176118

Selman, M., & Pardo, A. (2021). Fibroageing: An ageing pathological feature driven by dysregulated extracellular matrix-cell mechanobiology. Ageing Research Reviews, 70, 101393.

Simon, A., & Tanaka, E. M. (2013). Limb regeneration. Wiley Interdisciplinary Reviews. Developmental Biology, 2(2), 291–300.

Sousounis, K., Athippozhy, A. T., Voss, S. R., & Tsonis, P. A. (2014). Plasticity for axolotl lens regeneration is associated with age-related changes in gene expression. *Regeneration (Oxford*, England), 1(3), 47–57.

Sperber, H., Mathieu, J., Wang, Y., Ferreccio, A., Hesson, J., Xu, Z., Fischer, K. A., Devi, A., Detraux, D., Gu, H., Battle, S. L., Showalter, M., Valensisi, C., Bielas, J. H., Ericson, N. G., Margaretha, L., Robitaille, A. M., Margineantu, D., Fiehn, O., … Ruohola-Baker, H. (2015). The metabolome regulates the epigenetic landscape during naive-to-primed human embryonic stem cell transition. Nature Cell Biology, 17(12), 1523–1535.

Stocum, D. L. (2017). Mechanisms of urodele limb regeneration. *Regeneration (Oxford*, England), 4(4), 159–200.

Stocum, D. L., & Cameron, J. A. (2011). Looking proximally and distally: 100 years of limb regeneration and beyond. Developmental Dynamics: An Official Publication of the American Association of Anatomists, 240(5), 943–968.

Struck, A.-W., Thompson, M. L., Wong, L. S., & Micklefield, J. (2012). S-adenosyl-methionine-dependent methyltransferases: highly versatile enzymes in biocatalysis, biosynthesis and other biotechnological applications. Chembiochem: A European Journal of Chemical Biology, 13(18), 2642–2655.

Suetsugu-Maki, R., Maki, N., Nakamura, K., Sumanas, S., Zhu, J., Del Rio-Tsonis, K., & Tsonis, P. A. (2012). Lens regeneration in axolotl: new evidence of developmental plasticity. BMC Biology, 10, 103.

Suthon, S., Perkins, R. S., Bryja, V., Miranda-Carboni, G. A., & Krum, S. A. (2021). WNT5B in Physiology and Disease. Frontiers in Cell and Developmental Biology, 0. 10.3389/fcell.2021.667581

Sylva, M., Moorman, A. F. M., & van den Hoff, M. J. B. (2013). Follistatin-like 1 in vertebrate development. Birth Defects Research Part C: Embryo Today: Reviews, 99(1), 61–69.

Szklarczyk, D., Kirsch, R., Koutrouli, M., Nastou, K., Mehryary, F., Hachilif, R., Gable, A. L., Fang, T., Doncheva, N. T., Pyysalo, S., Bork, P., Jensen, L. J., & von Mering, C. (2023). The STRING database in 2023: protein-protein association networks and functional enrichment analyses for any sequenced genome of interest. Nucleic Acids Research, 51(D1), D638–D646.

Takeshita, S., Fumoto, T., Matsuoka, K., Park, K.-A., Aburatani, H., Kato, S., Ito, M., & Ikeda, K. (2013). Osteoclast-secreted CTHRC1 in the coupling of bone resorption to formation. The Journal of Clinical Investigation, 123(9), 3914–3924.

Tang, B. L. (2001). ADAMTS: a novel family of extracellular matrix proteases. The International Journal of Biochemistry & Cell Biology, 33(1), 33–44.

Tazaki, A., Tanaka, E. M., & Fei, J.-F. (2017). Salamander spinal cord regeneration: The ultimate positive control in vertebrate spinal cord regeneration. Developmental Biology, 432(1), 63–71.

UniProt Consortium. (2023). UniProt: the Universal Protein Knowledgebase in 2023. Nucleic Acids Research, 51(D1), D523–D531.

Vance, E. (2017). Biology’s beloved amphibian — the axolotl — is racing towards extinction. Nature, 551(7680), 286–289.

van Duyvenvoorde, H. A., Lui, J. C., Kant, S. G., Oostdijk, W., Gijsbers, A. C. J., Hoffer, M. J. V., Karperien, M., Walenkamp, M. J. E., Noordam, C., Voorhoeve, P. G., Mericq, V., Pereira, A. M., Claahsen-van de Grinten, H. L., van Gool, S. A., Breuning, M. H., Losekoot, M., Baron, J., Ruivenkamp, C. A. L., & Wit, J. M. (2014). Copy number variants in patients with short stature. European Journal of Human Genetics: EJHG, 22(5), 602–609.

Vieira, W. A., Wells, K. M., & McCusker, C. D. (2020). Advancements to the Axolotl Model for Regeneration and Aging. Gerontology, 66(3), 212–222.

Voss, S. R., Smith, J. J., Cecil, R. F., Kabangu, M., Duerr, T. J., Monaghan, J. R., Timoshevskaya, N., Ponomareva, L. V., Thorson, J. S., Veliz-Cuba, A., & Murrugarra, D. (2021). HDAC Inhibitor Titration of Transcription and Axolotl Tail Regeneration. Frontiers in Cell and Developmental Biology, 9, 767377.

Voss, S. R., Woodcock, M. R., & Zambrano, L. (2015). A Tale of Two Axolotls. Bioscience, 65(12), 1134–1140.

Whitcomb, S. J., Basu, A., Allis, C. D., & Bernstein, E. (2007). Polycomb Group proteins: an evolutionary perspective. Trends in Genetics: TIG, 23(10), 494–502.

Whited, J. L., Lehoczky, J. A., Austin, C. A., & Tabin, C. J. (2011). Dynamic expression of two thrombospondins during axolotl limb regeneration. Developmental Dynamics: An Official Publication of the American Association of Anatomists, 240(5), 1249–1258.

Wu, C.-H., Huang, T.-Y., Chen, B.-S., Chiou, L.-L., & Lee, H.-S. (2015). Long-duration muscle dedifferentiation during limb regeneration in axolotls. PloS One, 10(2), e0116068.

Xuan, F., Yano, F., Mori, D., Chijimatsu, R., Maenohara, Y., Nakamoto, H., Mori, Y., Makii, Y., Oichi, T., Taketo, M. M., Hojo, H., Ohba, S., Chung, U.-I., Tanaka, S., & Saito, T. (2019). Wnt/β-catenin signaling contributes to articular cartilage homeostasis through lubricin induction in the superficial zone. Arthritis Research & Therapy, 21(1), 247.

Yang, Y., Liu, J., Mao, H., Hu, Y.-A., Yan, Y., & Zhao, C. (2009). The expression pattern of Follistatin-like 1 in mouse central nervous system development. Gene Expression Patterns: GEP, 9(7), 532–540.

Yariv, B., Yariv, E., Kessel, A., Masrati, G., Chorin, A. B., Martz, E., Mayrose, I., Pupko, T., & Ben-Tal, N. (2023). Using evolutionary data to make sense of macromolecules with a “face-lifted” ConSurf. Protein Science: A Publication of the Protein Society, 32(3), e4582.

Yu, X., Kline, B., Han, Y., Gao, Y., Fan, Z., & Shi, Y. (2022). Weill-Marchesani syndrome 4 caused by compound heterozygosity of a maternal submicroscopic deletion and a paternal nonsense variant in the gene: A case report. American Journal of Ophthalmology Case Reports, 26, 101541.

Zhou, Q., Lyu, S., Bertrand, A. A., Hu, A. C., Chan, C. H., Ren, X., Dewey, M. J., Tiffany, A. S., Harley, B. A. C., & Lee, J. C. (2021). Stiffness of Nanoparticulate Mineralized Collagen Scaffolds Triggers Osteogenesis via Mechanotransduction and Canonical Wnt Signaling. Macromolecular Bioscience, 21(3), e2000370.

